# Allosteric mechanism of transcription inhibition by NusG-dependent pausing of RNA polymerase

**DOI:** 10.1101/2022.10.28.514324

**Authors:** Rishi K. Vishwakarma, M. Zuhaib Qayyum, Paul Babitzke, Katsuhiko S. Murakami

## Abstract

NusG is a transcription elongation factor that stimulates transcription pausing in Gram+ bacteria including *Bacillus subtilis* by sequence-specific interaction with a conserved pause-inducing _-11_TTNTTT_-6_ motif found in the non-template DNA (ntDNA) strand within the transcription bubble. To reveal the structural basis of NusG-dependent pausing, we determined a cryo-EM structure of a paused transcription complex containing RNAP, NusG, and the TTNTTT motif in the ntDNA strand. Interaction of NusG with the ntDNA strand rearranges the transcription bubble by positioning three consecutive T residues in a cleft between NusG and the β-lobe domain of RNAP. We revealed that the RNAP swivel module rotation (swiveling), which widens (swiveled state) and narrows (non-swiveled state) a cleft between NusG and the β-lobe, is an intrinsic motion of RNAP and is directly linked to nucleotide binding at the active site and to trigger loop folding, an essential conformational change of all cellular RNAPs for the RNA synthesis reaction. We also determined cryo-EM structures of RNAP escaping from a paused transcription complex. These structures revealed the NusG-dependent pausing mechanism by which NusG-ntDNA interaction inhibits the transition from swiveled to non-swiveled states, thereby preventing trigger loop folding and RNA synthesis allosterically. This motion is also reduced by formation of an RNA hairpin within the RNA exit channel. Thus, the pause half-life can be modulated by the strength of the NusG-ntDNA interaction and/or the stability of the RNA hairpin. NusG residues that interact with the TTNTTT motif are widely conserved in bacteria, suggesting that NusG-dependent pausing of transcription is widespread.

**Significance statement:** Transcription pausing by RNA polymerase (RNAP) regulates gene expression where it controls co-transcriptional RNA folding, synchronizes transcription with translation, and provides time for binding of regulatory factors. Transcription elongation factor NusG stimulates pausing in Gram+ bacteria including *Bacillus subtilis* and *Mycobacterium tuberculosis* by sequence-specific interaction with a conserved pause motif found in the non-template DNA (ntDNA) strand within the transcription bubble. Our structural and biochemical results revealed that part of the conserved TTNTTT motif in ntDNA is extruded and sandwiched between NusG and RNAP. Our results further demonstrate that an essential global conformational change in RNAP is directly linked to RNA synthesis, and that the NusG-ntDNA interaction pauses RNA synthesis by interfering with this conformational change.

## Introduction

Gene expression requires a multi-subunit DNA-dependent RNA polymerase (RNAP) in all organisms. Although RNA synthesis during transcription elongation by RNAP is highly processive, it is occasionally disrupted by random or programmed offline states, called transcription pausing. Temporal inactivation of RNA synthesis underlies diverse mechanisms that regulate gene expression from bacteria to human (1, 2). In bacteria, transcription pausing facilitates: 1) folding of nascent transcripts into structures that function as ribozymes and riboswitches (3–5); 2) coupling of transcription and translation (6–8); 3) provides time for binding of regulatory factors to the transcription elongation complex (TEC) or the nascent transcript (9–11); and 4) enables transcription termination (12–14).

Transcription pausing is triggered by interactions among RNAP and nucleic acid strands (RNA and DNA) with or without regulatory factors (15). Some of these interactions interrupt the nucleotide addition cycle by promoting entry of RNAP into a short-lived elemental paused state, that can lead to long-lived paused or backtracked states or to transcription termination (1, 2) Structural studies of the elemental paused complex using *Escherichia coli* RNAP revealed a half-translocated and tilted RNA-DNA hybrid (RNA post-translocated, DNA pre-translocated) that pauses transcription by impairing binding of an incoming nucleotide (NTP) at the active site of RNAP (16–18). A cryo-EM structure of an RNA hairpin-stabilized paused transcription complex (PTC) in which the RNA hairpin forms within the RNA exit channel not only forms a tilted RNA-DNA hybrid but also rotates a swivel module (ω subunit + clamp and shelf domains) of RNAP toward the upstream DNA (aka swiveled conformation), further stabilizing the paused transcription state by preventing folding of the trigger loop, an essential step for RNA synthesis (16, 17, 19). Trigger loop changes its folding depending on NTP binding at the active site. Without NTP, the trigger loop is unfolded and conserved residues such as M1012 and H1016 (*Mycobacterium tuberculosis* RNAP numbering) are 30 Å away from the NTP binding site. With correct NTP loading at the active site, the trigger loop folds as the two α-helices form extensive interactions that include contacts of M1012 and H1016 with the nucleobase and triphosphate group of the NTP, respectively. These interactions place the NTP into position for the nucleotidyl transfer reaction (i.e., at the insertion site) (20). Numerous studies established *E. coli* RNAP as a paradigm for elucidating fundamental mechanisms of transcription and its regulation of gene expression. However, transcription pausing with the *E. coli* transcription system links Si3, a 188 amino acid insertion domain found in the middle of the *E. coli* RNAP trigger loop (17, 21, 22). Yet, Si3 is a lineage-specific insertion found only in proteobacteria, and most other phyla including *M. tuberculosis* (*Mtb*) and *B. subtilis* RNAP lack this insertion. Thus, this transcription pausing mechanism is not widespread in bacteria.

Transcription elongation factors NusA and NusG influence transcription pausing *in vitro* and *in vivo* (2, 23). NusA stabilizes the paused state of transcription elongation by promoting the formation of an RNA hairpin (16). *E. coli* NusG and its paralog RfaH suppress RNAP backtracking by stabilizing the upstream duplex DNA and by preventing RNAP from forming the swiveled conformation (24, 25), thereby increasing transcription speed and processivity. RfaH establishes sequence-specific ntDNA contacts within the transcription bubble and this tight binding of RfaH to the transcription complex prevents RNA hairpin-stabilized pausing (9, 25–29).

In *B. subtilis*, interaction of NusG with the ntDNA strand stimulates transcriptional pausing (30). Genome-wide sequencing of nascent RNA followed by RNase I digestion (RNET-seq) revealed ∼1,600 strong NusG-dependent pause sites containing a conserved _-11_TTNTTT_-6_ motif in the ntDNA strand within the PTC (31). NusG-dependent pausing is enhanced by an RNA hairpin that forms ∼11-13 nt upstream of the 3’ end of the paused transcript but the RNA hairpin is not required for pausing (2). NusG makes sequence-specific contacts with the TTNTTT motif; NusG deletion or point mutations in NusG or the TTNTTT motif greatly reduces pausing. Four NusG-dependent pause sites in 5’UTRs (*trpE, tlrB, ribD,* and *vmlR*) have been characterized and shown to be involved in regulating expression of the downstream gene (2, 11, 32, 33). In addition, pausing plays an important role in NusG-stimulated intrinsic termination; deletion of NusG results in the misregulation of global gene expression and altered cellular physiology (14). Phylogenetic analysis of NusG in bacteria revealed that the *B. subtilis* type of NusG is widespread but not in proteobacteria (31). Moreover, it was shown that Spt5, the NusG homolog in eukaryotes, also contacts the ntDNA strand within the transcription bubble, which may regulate transcription (34). These findings suggest that NusG-ntDNA interaction within the transcription bubble might be a universally conserved mechanism of transcription pausing in all organisms.

In this study, we investigated the mechanism of NusG-dependent pausing by determining a series of cryo-EM structures of transcription complexes containing *Mtb* RNAP, DNA/RNA nucleic acid scaffolds, and NusG from *B. subtilis* or *Mtb*. These complexes include elongation and paused transcription states, as well as complexes undergoing pause escape. We revealed two functions of the TTNTTT motif and a role of the RNA hairpin in pausing and in determining the pause half-life, which is a measure of pause duration. We also revealed a direct link between global (swiveling) and local (trigger loop folding) conformational changes of RNAP during the nucleotide addition cycle, and that NusG-dependent pausing prevents RNA synthesis allosterically by interfering with rotation of the RNAP swivel module.

## Results

### B. subtilis NusG induces pausing of Mtb and B. subtilis RNAPs in response to a conserved TTNTTT motif

*B. subtilis* NusG stimulates pausing of RNAP via interaction of its N-terminal NGN domain with a conserved TTNTTT motif in the ntDNA strand within the transcription bubble (30, 31). During our cryo-EM studies on transcription, we found that *B. subtilis* RNAP is poorly suited for cryo-EM structural studies due to its weak association with the nucleic acid scaffold and dissociation of RNAP subunits from the complex during sample vitrification. We therefore tested *Mtb* RNAP, which has been used for cryo-EM structural studies and is closely related to *B. subtilis* RNAP. To evaluate whether *Mtb* RNAP transcription pausing is stimulated by *B. subtilis* NusG, we performed *in vitro* transcription assays using these RNAPs and a strong NusG-dependent pause site within the *coaA* coding sequence **(Fig. 1A**). *B. subtilis* NusG stimulated pausing of both RNAPs at the same position and to similar extents. NusG increased the pause half-life of both RNAPs ∼4-5 fold (**Fig. 1B**). When the TTTTTT sequence was replaced with AACAAA, NusG-dependent pausing of both RNAPs was eliminated (**Fig. 1C**). These results established that *B. subtilis* NusG stimulates pausing of both RNAPs at the same position in response to the TTNTTT motif, indicating that *Mtb* RNAP can be used as a model for investigating the structural basis of NusG-dependent pausing.

**Figure 1:**
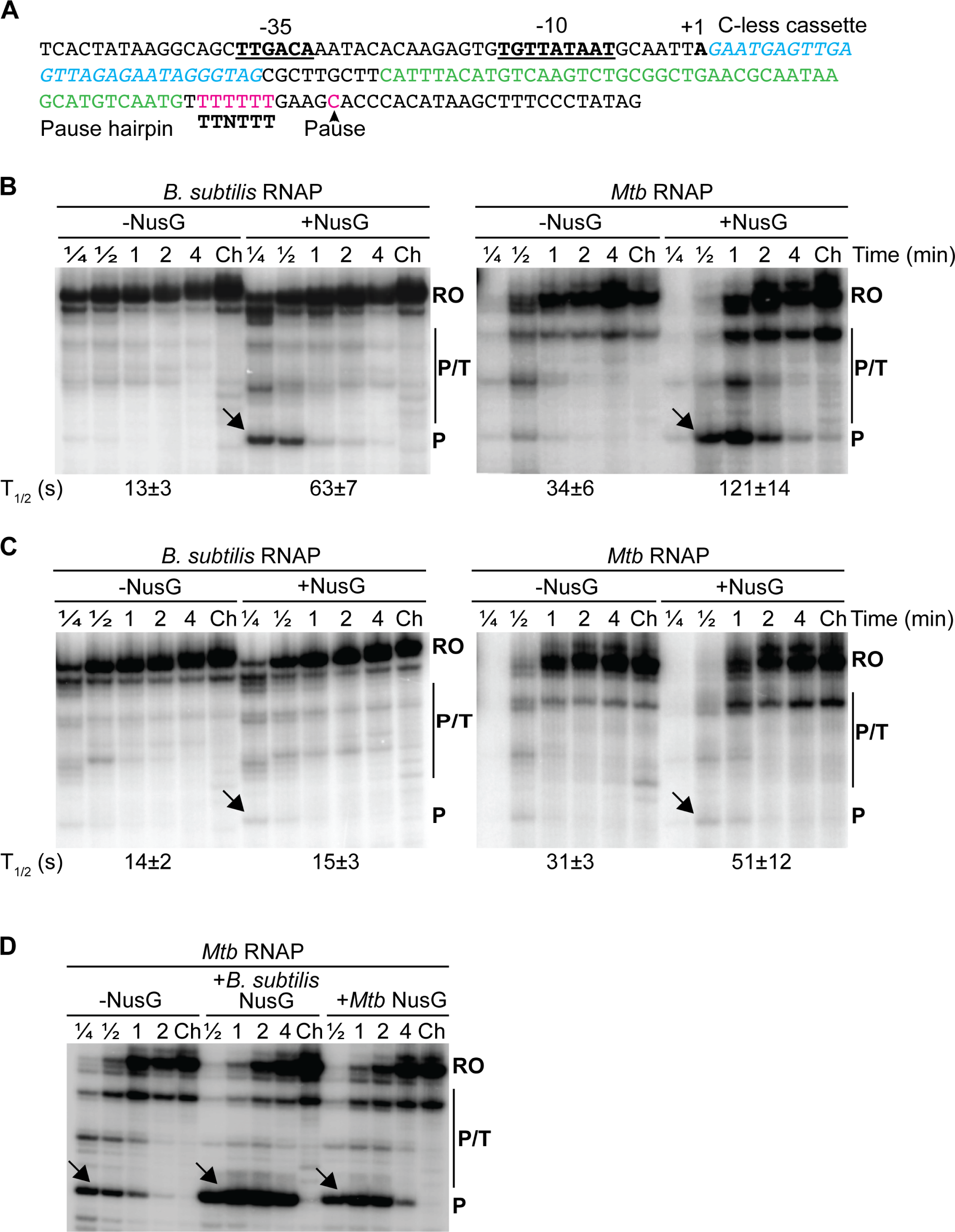
*B. subtilis* NusG stimulates TTNTTT motif-specific pausing of *Mtb* and *B. subtilis* RNAPs. **A.** DNA sequence used for promoter-initiated *in vitro* pausing assays. The -35 and extended -10 promoter elements are underlined. A 29-nucleotides C-less region is in cyan and the predicted *coaA* pause hairpin is in green. The TTNTTT motif and the pause position (arrowhead) are in magenta. The transcription start site (+1) is also indicated. **B.**Representative gels showing RNAs generated from promoter-initiated single-round *in vitro* transcription pausing assays. Reactions were performed with the *coaA* DNA template shown in (**A**), *B. subtilis* NusG and either *B. subtilis* (left) or *Mtb* (right) RNAP. Transcription reactions were performed with 150 μM of each NTP ± 1 μM NusG. Reactions were stopped at the times shown above each lane. Chase reactions (Ch) were extended for an additional 10 min at 37 °C. Positions of the *coaA* pause (P and arrow), and run-off (RO) transcripts are indicated. Additional pause, terminated, or arrested RNA species (P/T) were observed between P and RO. Calculated pause half-lives are shown at the bottom of the gel. **C.** Representative gels showing RNAs from the promoter-initiated single-round *in vitro* transcription reactions using a DNA template in which the TTNTTT motif was replaced with AACAAA. Values at the bottom of the gels are averages ± standard deviation (n=3). **D**. *Mtb* NusG stimulates pausing of *Mtb* RNAP at the same position as *B. subtilis* NusG. Reactions were performed in the absence (-NusG) and in the presence of either *B. subtilis* or *Mtb* NusG as indicated. Transcription reactions were performed with 150 μM of each NTP except ATP (10 µM). The DNA template used for this analysis contained the RNA hairpin sequence used for reconstruction of the PTC and TEC (Fig. 2A) rather than the natural *coaA* hairpin sequence (green in panel A).

### Preparation of a NusG-dependent PTC using a nucleic acid scaffold containing the TTNTTT motif

TECs have been conveniently prepared using nucleic acid scaffolds for *in vitro* transcription assays and for structural studies (17, 30, 35–37). We tested RNA extension and pause escape of *Mtb* RNAP using a nucleic acid scaffold containing the _-11_TTNTTT_-6_ motif relative to the pause position at -1, a 30-mer RNA with an RNA hairpin (7 bp stem and 4 base loop) at its 5’ end, and 9 bases complementary to the template DNA (tDNA) strand (**Fig. 2A**). As for promoter-derived transcription complexes (**Fig. 1B**), RNA extension (from 30 to 33-mer in the presence of ATP, GTP and 3’-dCTP) was delayed in response to both NusG and the TTNTTT motif (**Fig. 2B** and **2C**). These results established that NusG-dependent PTCs can be prepared using *Mtb* RNAP, nucleic acid scaffolds and *B. subtilis* NusG for structure determination.

**Figure 2:**
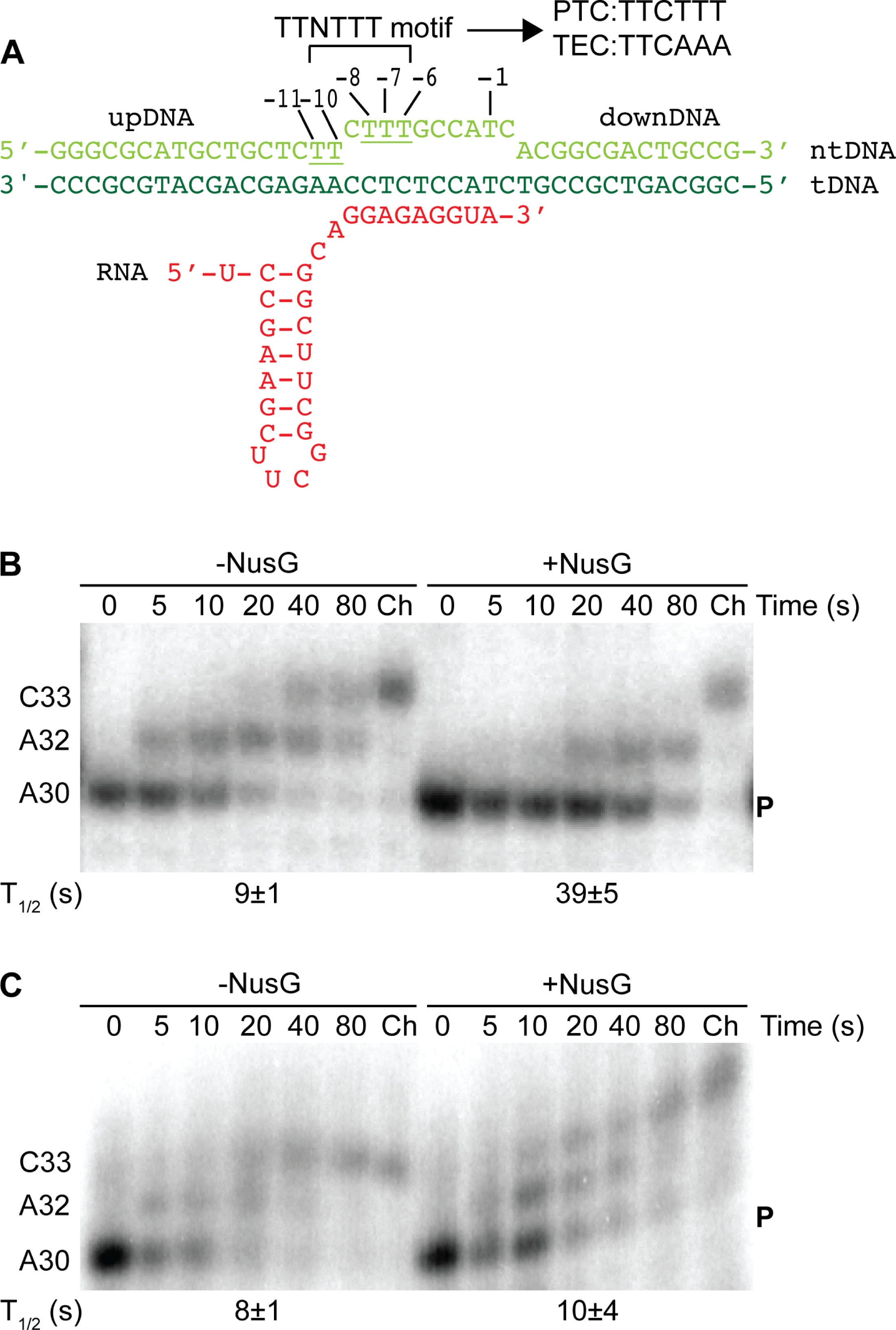
Reconstitution of active PTCs with a DNA/RNA scaffold, *Mtb* RNAP and *B. subtilis* NusG. **A.** Nucleic acid scaffold used for reconstitution of the active PTC and TEC. Non-template DNA (ntDNA) for preparing the PTC contained the TTNTTT motif (underlined). The last three T residues of the TTNTTT motif (positions -8 to -6) were replaced with A residues (TTCAAA) for preparing the TEC. upDNA, upstream DNA; downDNA, downstream DNA. **B.** Representative gels showing the results of single-round *in vitro* pause escape assays using the PTC nucleic acid scaffold. Reactions were stopped at the times shown above each lane. Chase reactions (Ch) were extended for an additional 10 min at 37 °C. **C**. Same as panel (**B**) except the nucleic acid scaffold for the TEC was used. Values at the bottom of the gels are averages ± standard deviation (n=3).

### Cryo-EM structures of TEC and NusG-dependent PTC

We prepared PTCs and TECs using nucleic acid scaffolds with and without the TTNTTT motif (**Fig. 2A**) and determined cryo-EM structures for both complexes to a nominal resolution of about 3 Å (**Figs. S1, S2 and Table S1**). The issue of preferred particle orientation, a highly biased particle orientation due to particles facing air within a thin layer of water, during sample vitrification was resolved by adding 8.0 mM (final concentration) of CHAPSO to the sample just before applying to the cryo-EM grid (38). Both transcription complexes show well-defined densities for RNAP, the NGN domain of NusG, and the nucleic acid scaffold except for RNA upstream of the RNA-DNA hybrid, which is relatively weak but still sufficient to position the RNA hairpin model (**Figs. 3A, 3B, S3A and Movie S1**).

**Figure 3:**
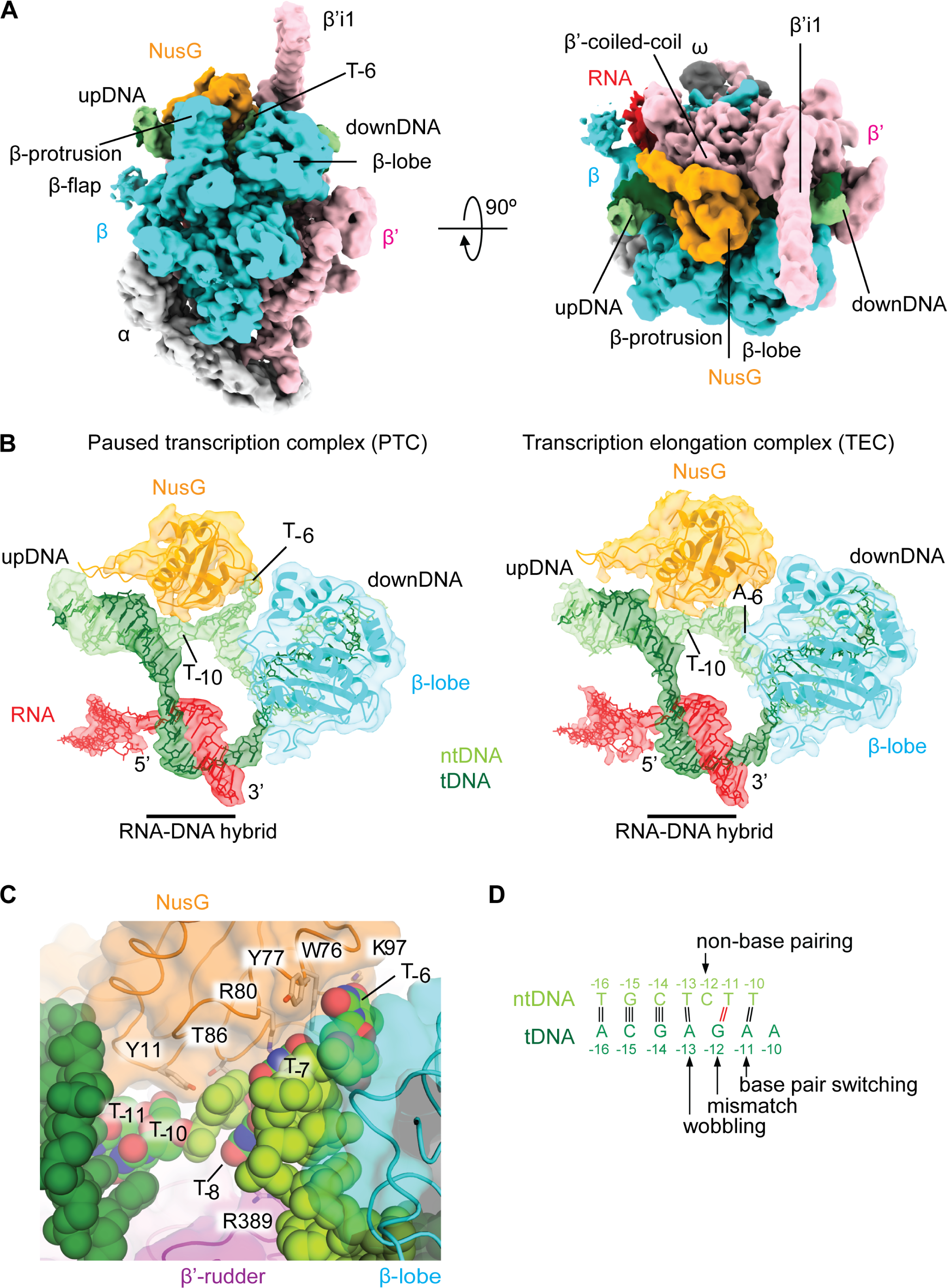
Cryo-EM structures of the TEC and NusG-dependent PTC. **A.** Orthogonal views of the cryo-EM density map of the PTC. Subunits and domains of RNAP, DNA, RNA and NusG are colored and labeled. downDNA, downstream DNA; upDNA, upstream DNA; β’-i1, lineage-specific insertion. **B.** Cryo-EM densities (transparent) of DNA, RNA, β-lobe domain and NusG are overlayed with the PTC structure (left) and TEC structure (right) revealing the rearrangement of the transcription bubble preceding the upstream DNA in the PTC. The 5’ and 3’ ends of the RNA, as well as positions -10 and -6 of the ntDNA are indicated. The cryo-EM density maps and the structure are colored according to (**A**). **C.** A magnified view of the ntDNA interactions with NusG and RNAP (same orientation as in the panel B). Amino acid residues of NusG and the β’ subunit interacting with ntDNA are indicated as stick models. DNA strands (tDNA, dark green; ntDNA, light green) are shown as sphere models and thymine residues in the TTNTTT motif are colored (red, oxygen; blue, nitrogen; green, carbon). **D**. Schematic representations of the base pairing distortion of the upstream DNA

The density of the ntDNA in the PTC is sufficiently clear for modeling all bases in the single-stranded DNA within the transcription bubble. In contrast, the ntDNA density in the TEC without the TTNTTT motif is relatively weak, suggesting that it is flexible. The C-terminal domain of the α subunit (α-CTD) and the C-terminal domain of NusG (KOW domain) are visible in the low path filtered map (**Fig. S3B**). The NGN domain of NusG binds two pincers of the RNAP main channel including the coiled-coil helices of the β’ clamp and the protrusion domain of the β subunit (β-protrusion) as its primary and secondary binding sites, respectively. This interaction fills a cleft of RNAP and covers the single-stranded ntDNA and the upstream double-stranded DNA (**Fig. 3B**). In both transcription complexes: 1) the density corresponding to the RNA-DNA hybrid is strong such that the 9 bp RNA-DNA hybrid is modeled without any ambiguity; 2) the 3’ end of the RNA is in the post-translocated site (*i* site) as observed in the genome-wide study of NusG-dependent pausing in *B. subtilis* (31); 3) the RNAP swivel module with NusG rotates ∼2.1 degrees toward the upstream DNA compared with the one in the escaped TEC with NTP as described later, and 4) the RNA-DNA hybrid is not in a half-translocated tilted state (RNA post-translocated, DNA pre-translocated) as reported and used to explain the mechanism of transcription pausing for *E. coli* RNAP (**Fig. S3D**) (16, 17). These observations imply that the mechanism of NusG-dependent pausing is distinct from the mechanism reported for *E. coli* RNAP.

To explore the possibility that the cryo-EM structure may contain hidden conformation(s) of RNAP, NusG and/or the RNA-DNA hybrid, we analyzed the conformational heterogeneity of the cryo-EM datasets by performing global 3D variability analysis (3DVA) in cryoSPARC-v3.3 (39, 40). We did not observe conformational heterogeneity of RNAP or the RNA-DNA hybrid in the PTC data set; however, we found that one of the variability components in the TEC showed: 1) swivel module rotation relative to the core module; 2) movement of the upstream DNA; and changes in the size of the RNA exit channel (**Movies S2 and S3**). Particles were sorted into two clusters along this 3D conformational trajectory followed by 3D refinement, which yielded two structure classes (class 1, 48% of the particles; class 2, 52% of the particles; **Fig. S2**). Comparison of the class 1 and class 2 structures revealed that the RNAP swivel module with NusG rotates ∼1.5 degrees toward the downstream DNA, with the rotation axis positioned at the ω subunit that roughly parallels the bridge helix (**Fig. 4 A and B**). This conformational change of RNAP, including the moving swivel module and the rotation axis, is the same as reported in previous structural studies of *E. coli* RNAP (16, 17, 41) (**Movie S5**). We refer to these classes as the swiveled (class 1) and non-swiveled (class 2) conformations. RNAP swiveling changes the width of a cleft between NusG and the β-lobe domain and the size of the RNA exit channel (both are wider in the swiveled conformation and narrower in the non-swiveled conformation) (**Figs. 4B**, **S4**, and **Movie S3**). In the non-swiveled conformation of the TEC, the RNA exit channel is too narrow to accommodate an RNA hairpin and T_-6_ of ntDNA and hence the RNA hairpin was unfolded (**Fig. 4B right**). In the TEC, RNAP formed both the swiveled and non-swiveled conformations and the RNA hairpin was only observed in the swiveled conformation, suggesting that the RNA hairpin promotes formation of the swiveled conformation.

**Figure 4:**
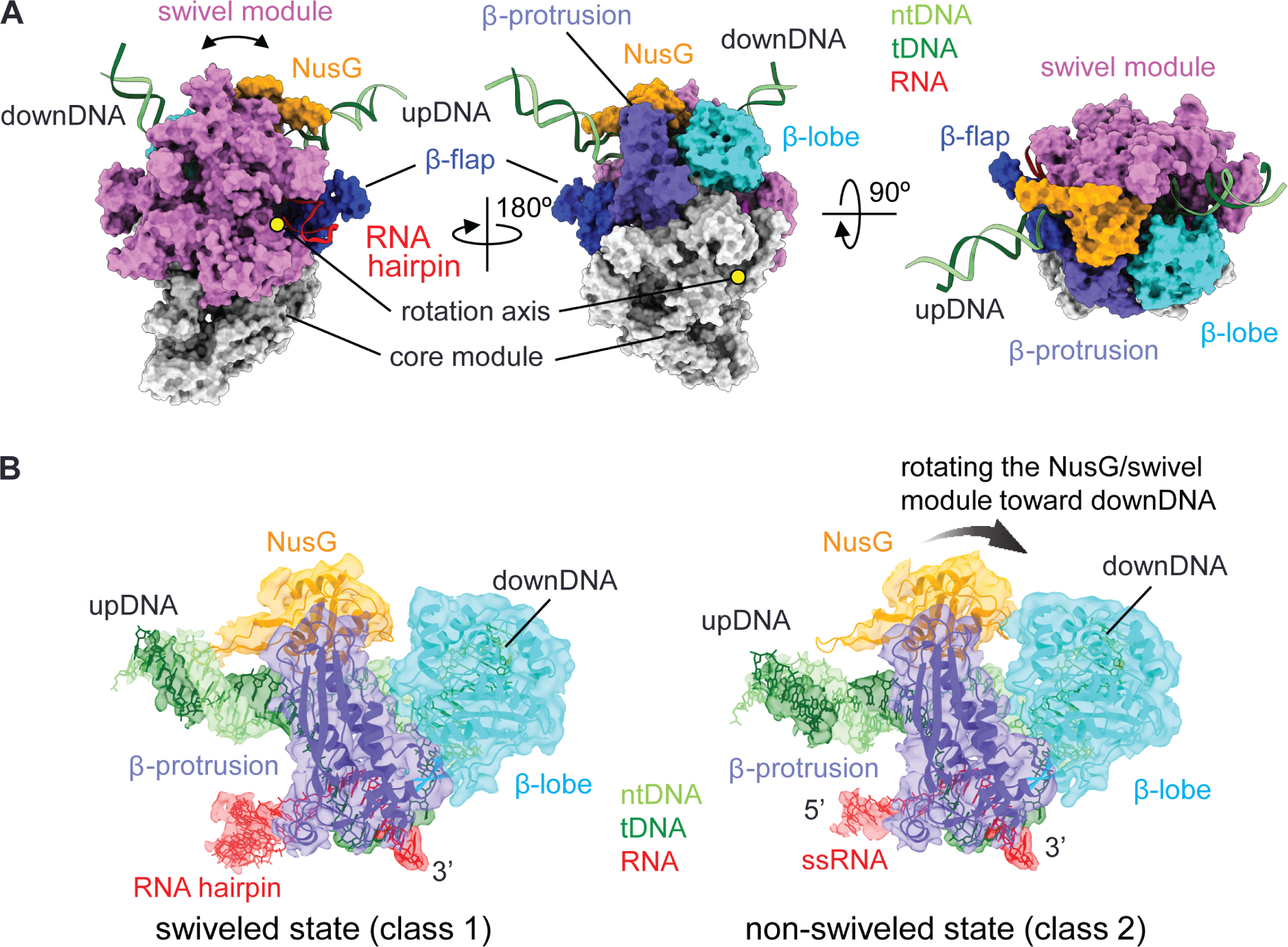
RNAP swiveling of *Mtb* RNAP. **A.** Structure of the *Mtb* RNAP-*B. subtilis* NusG TEC indicating the modules and domains. The rotation axis of the swivel module is shown as yellow circles (left and middle). **B.** Cryo-EM densities (transparent) of DNA, RNA, NusG, β-lobe and β-protrusion domains are overlayed with the swiveled (left) and non-swiveled states (right) of the TEC. Rotation of the NusG/swivel module from the swiveled to non-swiveled states is indicated by a black arrow (right panel).

### NusG interaction with the ntDNA strand rearranges the transcription bubble of the PTC

In the TEC, the ntDNA winds past NusG and the β-lobe domain without any interaction. In sharp contrast, the ntDNA is buried in a narrow cleft between NusG and the β-lobe domain in the PTC, and all residues in the TTNTTT motif interact with NusG and/or RNAP (**Fig. 3C**). Y11 of NusG contacts the T_-10_ backbone and C_-9_. In addition, T86 contacts C_-9_, while T_-8_ forms base stacking interactions with C_-9_ and R389 of the β’ subunit, which is positioned in the middle of the β’-rudder. T_-7_ also stacks with G_-5_ and with R80 and N81 of NusG. Furthermore, T_-6_ flips out from the ntDNA strand and fits snugly into a cavity of NusG formed by W76, Y77 and K97, and establishes van der Waals interactions with W76 and Y77. The position of T_-6_ is further stabilized by a salt bridge with K97 of NusG. Alanine substitutions at these residues (W76A, Y77A, R80A, N81A, T86A, K97A and Y11A) reduce the pause half-life and pausing efficiency except for Y77A, which slightly increases the pause half-life but reduces the pausing efficiency, a measure of the fraction of RNAPs that pause at the pause site. (**Fig. S5**). Only a few base-specific interactions between the T residues (T_-7_ and T_-6_) and RNAP/NusG were observed, suggesting that the presence of the T residues in the TTNTTT motif is important for fitting small hydrophobic pyrimidine bases into a narrow hydrophobic cavity between NusG and the β-lobe instead of establishing extensive base-specific hydrogen bonds with RNAP and NusG (**Fig. 3C**).

Our structure-based biochemical assays identified key pause-promoting residues of *B. subtilis* NusG that support the prediction of bacterial lineages that might carry out NusG-dependent pausing. Next, we analyzed to what extent NusG residues that bind the ntDNA strand are conserved by performing a multiple sequence alignment of NusG from a wide variety of bacterial species. We also included RfaH, an *E. coli* NusG paralog, in this analysis (**Fig. S6**). Most of the residues involved in the NusG-ntDNA interaction are highly conserved in firmicutes, actinobacteria, deinococcus and cyanobacteria. These residues are not conserved in RfaH. NusG residues W76, R80 and K97 are highly conserved throughout the bacterial kingdom, while Y11 is moderately conserved with similar amino acid residues replacing them in some γ-proteobacteria. Among all NusG variants containing alanine substitutions that were tested, substitutions in Y11, W76 and R80 drastically reduced NusG-dependent pausing (**Fig. S5**), establishing the importance of these residues in pausing, and the likely existence of a common pausing mechanism in bacteria. We also tested *Mtb* NusG and found that it stimulates pausing of *Mtb* RNAP transcription at the same position as *B. subtilis* NusG (**Fig. 1D**).

### DNA base pair partner switching of TT residues results in a distortion of the upstream DNA

The DNA sequence logo of NusG-dependent pause sites in the *B. subtilis* genome contains two consecutive T residues corresponding to the -11 and -10 positions in the ntDNA (TTNTTT, underlined) (2). The mutation of T_-10_ to C resulted in ∼6-fold decreased pause half-life, whereas changing both T_-11_ and T_-10_ to Cs, resulted in ∼12-fold decreased pause half-life (42). The finding that mutations in these two residues did not eliminate pausing suggests that their role is to enhance the effect of the interaction between NusG and the three consecutive T residues at positions T_-8_, T_-7_ and T_-6_ in the transcription bubble. Although the backbone of T_-11_ and T_-10_ contact NusG, there is no base-specific interaction of these residues with NusG (**Fig. 3C**), raising the question as to how these two T residues contribute to NusG-dependent pausing. In the TEC, T_-11_ and T_-10_ in the ntDNA base pair with A_-11_ and A_-10_ in the tDNA as expected, resulting in B-form double-stranded DNA. In contrast, double-stranded DNA in the PTC is distorted because of DNA base pair switching, mismatch and wobbling (**Fig. 3B**). In this case, T_-_

_10_ (ntDNA) forms a base-pair with A_-11_ (tDNA) instead of A_-10_ (tDNA) such that A_-10_ remains single-stranded at the edge of the upstream DNA duplex (**Fig. 3D**). The base pair switching triggers a mismatch between T_-11_ (ntDNA) and G_-12_ (tDNA), no base pairing of C_-12_ (ntDNA), and base pair wobbling of T_-13_ (ntDNA) and A_-13_ (tDNA) (the distance between the N3 of T_-13_ and N1 of A_-13_ atoms is increased from 2.7 Å in the TEC to 4.4 Å in the PTC). Presumably, the presence of two consecutive thymine bases at the edge of the upstream DNA makes it prone to DNA base pair switching, thereby weakening closing of the transcription bubble during forward translocation of RNAP. Alternatively, the upstream DNA distortion facilitates rearrangement of the transcription bubble, which may establish stronger ntDNA interaction with NusG and the β-lobe (**Fig. 3A**). Elucidating the function of the upstream T residues of the TTNTTT motif during NusG-dependent pausing may require systematic investigation of this motif both *in vitro* (biochemistry and structural studies) and *in vivo*.

### Escape of RNAP from NusG-dependent pausing

Comparison of the *Mtb* RNAP structure conformations showed that RNAP was in the swiveled state in the PTC, whereas it shifts conformation between the swiveled and the non-swiveled state in the TEC. Therefore, we hypothesized that the NusG-ntDNA interaction may provide an obstacle to the rotation of the RNAP swivel module, and thereby cause RNA synthesis to pause. To test this hypothesis, we determined a series of cryo-EM structures that allowed us to directly observe RNAP escape from the PTC by mixing GMPCPP, a nonhydrolyzable analogue of GTP, to the PTC (**Figs. 5 and S7, Table S1**). The conformational heterogeneity of the complex was addressed by using the 3DVA in cryoSPARC-v3.3 (40). The analysis revealed a significant motion of the swivel module relative to the core module of RNAP, and the 3D refinement revealed three structure classes (**Fig. 5**). The first class does not show any density of the NTP at the active site and the RNAP/NusG/nucleic acid scaffold conformation is identical to the PTC (**Fig. 3**), thus we refer it as the PTC (26% of particles) (**Fig. 5B left**). The second class shows density of the NTP at the active site, but the RNAP/NusG/nucleic acid scaffold conformation is nearly identical to the PTC except for a slight rotation of the NusG/swivel module toward the downstream DNA and partial folding of the trigger loop. We refer to this structure as the PTC^‡^ + NTP (36% of particles) (**Fig. 5B middle**). The third class shows strong density of the NTP at the active site and the RNAP/NusG/nucleic acid scaffold conformation is the same as the TEC + NTP. We refer to this structure as the escaped transcription elongation complex eTEC + NTP (38% of particles) (**Fig. 5B right**). In the of PTC and PTC^‡^ + NTP structures, the last three T residues of the TTNTTT pause motif in ntDNA remain positioned in the narrow cavity between NusG and the β-lobe domain, and the RNA hairpin is folded. In the PTC^‡^ + NTP structure, full rotation of the RNAP swivel module toward downstream DNA is prohibited because of ntDNA binding within the NusG/β-lobe cavity. The trigger loop is partially folded in this structure such that M1012 contacts the base of the NTP, but H1016 cannot reach the triphosphate (**Fig. 5B** and **C**). In the eTEC + NTP structure: 1) the three T residues of the TTNTTT motif are pushed out of the NusG/β-lobe cavity, allowing full rotation of the swivel module to the non-swiveled state; 2) the trigger loop is fully folded to allow interaction of M1012 and H1016 with the NTP; 3) the second Mg^2+^ is loaded at the Mg^B^ site; and 1. 3) the RNA hairpin is unfolded (**Fig. 5 B and C**). In all three classes, the RNA-DNA hybrid configurations are identical without any evidence of tilting, and the RNAP 3’ end is in the post-translocated state. These observations indicate that NusG-dependent pausing does not interfere the NTP binding at the active site, but instead inhibits rotation of the RNAP swivel module by inserting ntDNA into the NusG/β-lobe cavity, thereby preventing completion of trigger loop folding, and hence causing RNA synthesis to pause.

**Figure 5:**
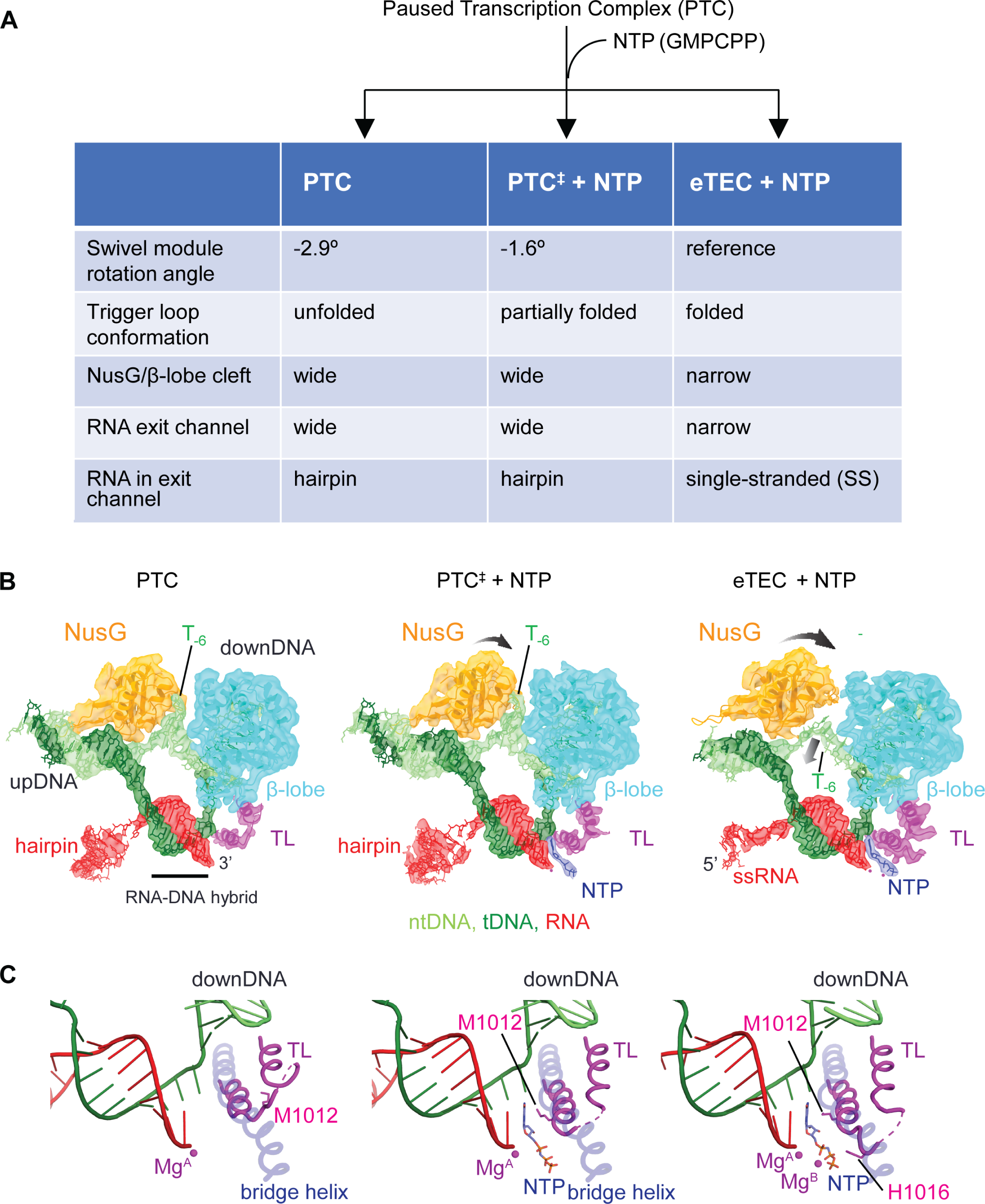
Pause escape from the PTC. **A.** Experimental scheme of the transcription complex preparation from the NusG-dependent PTC mixed with the incoming NTP. Three transcription complexes were formed, the angles of the swivel module relative to the eTEC + NTP are shown, and the conformations of the trigger loop are indicated. **B.** Comparison of the three transcription complexes. Cryo-EM densities (transparent) of DNA, RNA, NusG, β-lobe and trigger loop (TL) are overlayed with the final model. Rotation of NusG with the swivel module toward the downstream DNA are indicated by black arrows. Pushing the ntDNA out of the NusG/β-lobe cavity in the eTEC + NTP is indicated by a black arrow. **C.** Comparison of the trigger loop conformations in the three transcription complexes. Mg^2+^ ions at the active centers are shown as red spheres (Mg^A^ and Mg^B^), while the trigger loop residues M1012 and H1016 are shown as stick models.

### RNAP swiveling is an intrinsic motion of Mtb RNAP during the nucleotide addition cycle

From the structural study of the NusG-dependent PTC (**Fig. 5**), we found that the rotation of the RNAP swivel module is directly linked to the conformational change of the trigger loop, and the NusG-ntDNA interaction interferes with rotation of the RNAP swivel module thus inhibiting trigger loop folding and RNA synthesis. Next, to address whether RNAP swiveling is an intrinsic motion of *Mtb* RNAP during the nucleotide addition cycle, we prepared an *Mtb* RNAP TEC with *Mtb* NusG and a nucleic acid scaffold lacking the RNA hairpin and determined the cryo-EM structures (**Figs. 6, S8 and Table S1**). The structures of *Mtb* and *B. subtilis* NusG are nearly identical (RMSD=0.966 Å) except that *Mtb* NusG contains an extra α helix on its N-terminus (**Fig. 6B**). The amino acid residues of *Mtb* NusG facing the β-lobe domain, which are used for interacting with the ntDNA bases during NusG-dependent PTC formation, are well conserved with *B. subtilis* NusG, indicating that *Mtb* NusG may interact with the ntDNA bases to carry out NusG-dependent pausing (**Figs. 1D** and **6B**).

**Figure 6:**
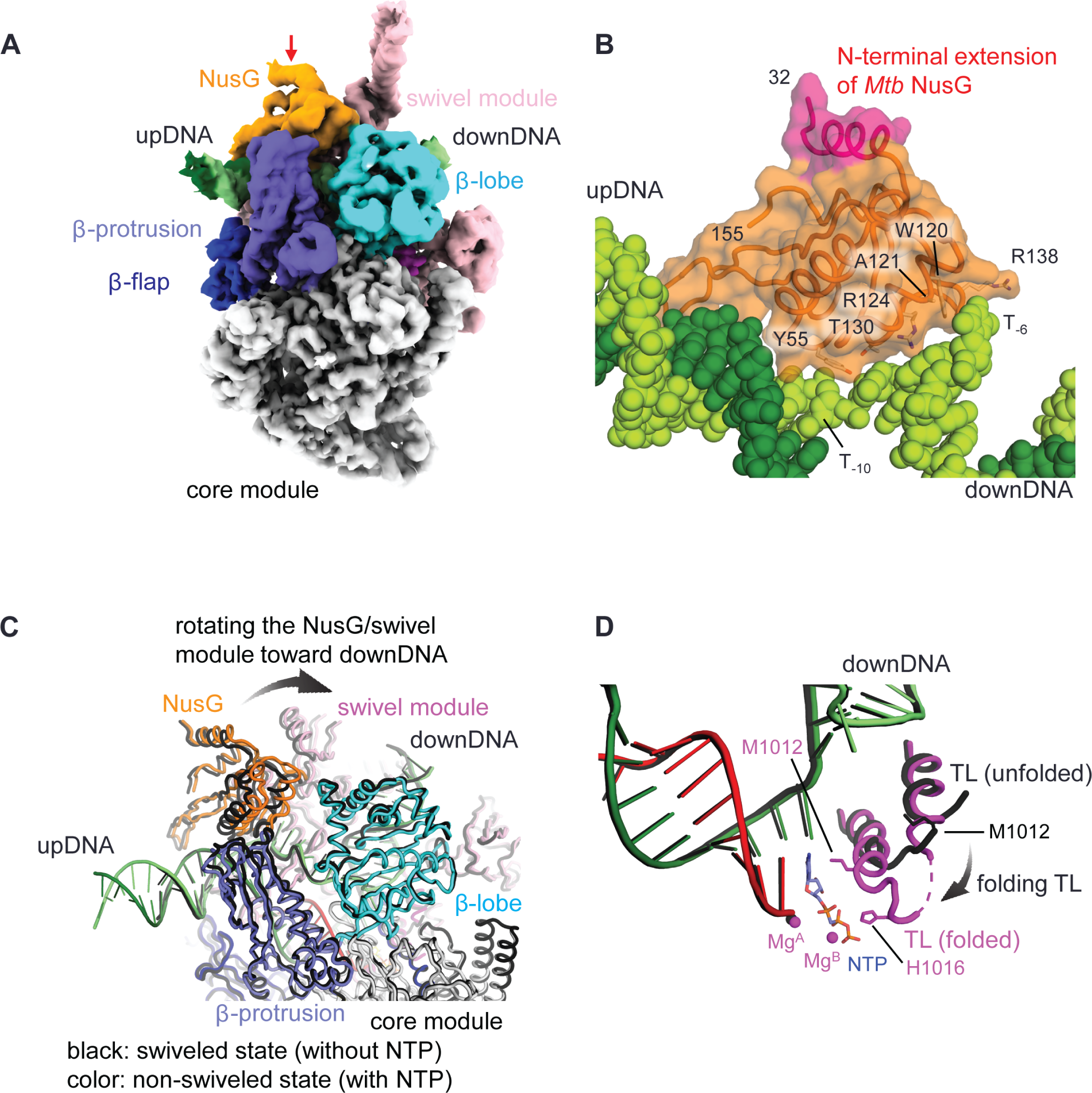
RNAP swiveling and trigger loop conformational changes of *Mtb* RNAP during the nucleotide addition cycle. **A.** The cryo-EM density map of the TEC containing *Mtb* RNAP and *Mtb* NusG with NTP. Modules and domains of RNAP, DNA, RNA and NusG are colored and labeled. The PTC is modeled by superposing *Mtb* NusG on *B. subtilis* NusG of the PTC structure. *Mtb* NusG *Mtb* NusG is indicated in red. **C,D**. Comparison of the structures of the *Mtb* RNAP-*Mtb* NusG TEC in the swiveled (without NTP, black) and non-swiveled (with NTP, color) states, showing the rotation of the NusG/swivel module (**C**) and folding of the trigger loop (**D**) upon NTP binding at the active site (indicated by black arrows). Mg^2+^ ions at active centers are shown as red spheres (Mg^A^ and Mg^B^) and the trigger loop residues M1012 and H1016 are shown as stick models that contact the nucleobase and triphosphate group of the NTP, respectively.

**Figure 7:**
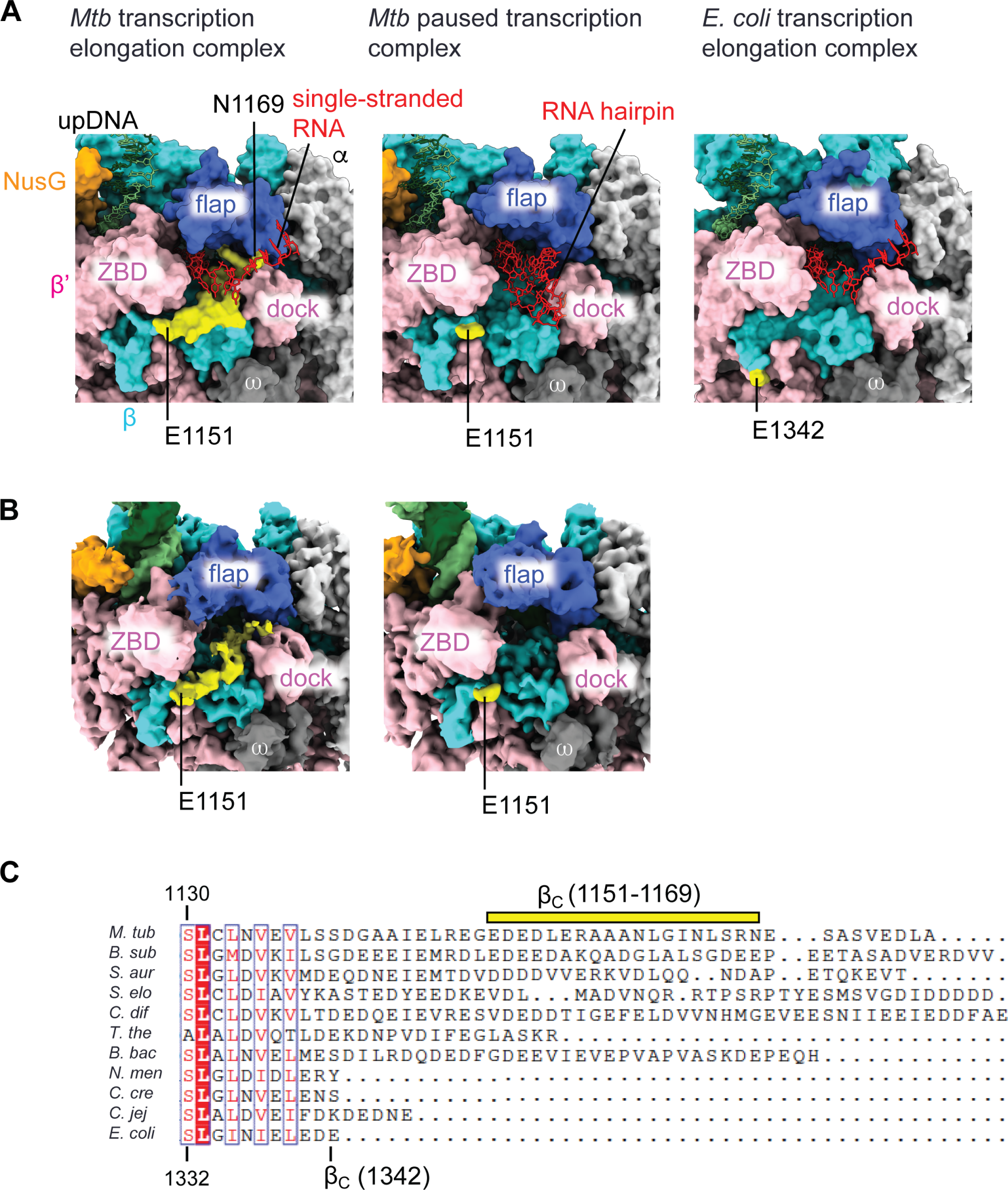
RNA exit channels of *Mtb* and *E. coli* RNAPs. **A.** Comparison of the RNA exit channels of the *Mtb* TEC without an RNA hairpin (left), the *Mtb* PTC with an RNA hairpin (middle), and the *E. coli* TEC without an RNA hairpin (right). The structures are depicted as surface representations (RNAP and NusG) and stick (DNA and RNA) models. The C-terminus of the *E. coli* β subunit (E1342, yellow) is positioned below the Zn binding domain (ZBD) of the β’ subunit (right). The C-terminal region of the *Mtb* β subunit (E1151 to N1169, yellow) forms a bridge between the Zn binding and dock domains of the β’ subunit, making the RNA exit channel narrower (left), but it is disordered in the PTC to accommodate the RNA hairpin (middle). **B.** Cryo-EM density maps of the TEC (left) and PTC (right). Densities of RNA are omitted. **C**. Sequence alignment of the C-terminus of the β subunit (βC) of representative bacterial RNAPs (*M. tub, M. tuberculosis; B. sub*, *B. subtilis*; *S. aur, Staphylococcus aureus*; *S. elo*, *Synechococcus elongatus*; *C. dif, Clostridioides difficile*; *T. the*, *Thermus thermophilus*; *B. bac, Bdellovibrio bacteriovorus*; *N. men, Neisseria meningitidis C*. *cre; Caulobacter crescentus*. *C. jej, Campylobacter jejuni*). The C-terminal tail of the *Mtb* β subunit and C-terminus of the *E. coli* β subunit are highlighted.

**Figure 8:**
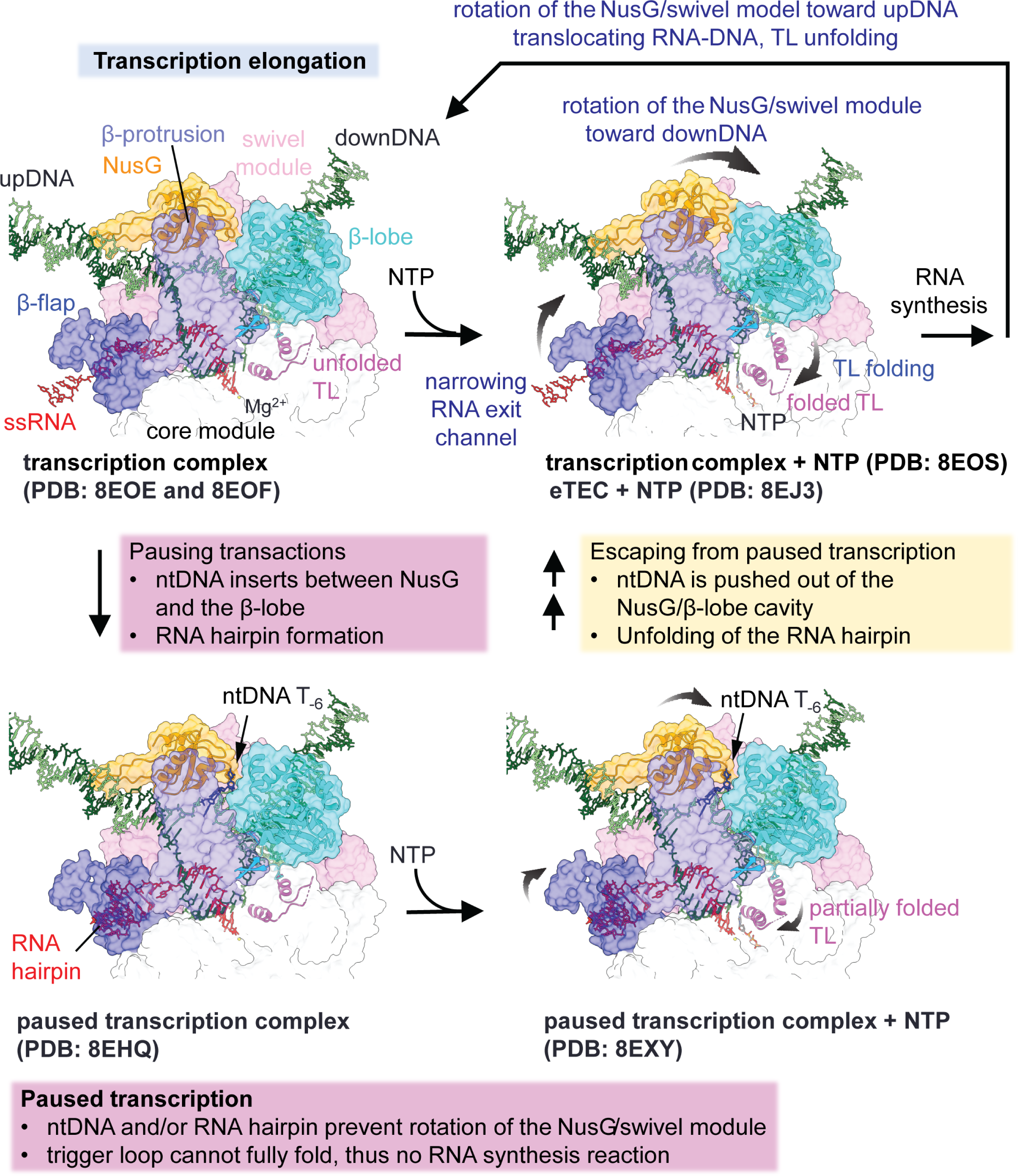
RNAP conformations associated with transcription elongation, NusG-dependent pausing, and escape from the PTC. A series of transcription complexes with NusG determined in this study are depicted as surface, ribbon and stick model representations, and the modules and domains are indicated. Top panels represent the TEC, while the bottom panels represent the PTC (left) and a complex escaping from the pause (eTEC) (right).

Next, we added CMPCPP, a non-hydrolyzable analogue of CTP, as an incoming NTP to monitor how binding of the incoming NTP at the RNAP active site influences the global (swiveling) and local (trigger loop folding) conformational changes of RNAP (**Fig. S9**). CryoSPARC-v3.3 3DVA analysis (40) revealed significant motion of the swivel module relative to the core module in the TECs, both in the presence and absence of NTP (**Movie S4**). In addition, sorting particles followed by 3D refinement revealed one major conformation in each TEC; swiveled without NTP and non-swiveled with NTP (**Fig. 6C**). The trigger loop is folded when NTP is bound at the active site (**Fig. 6D**). These results indicate that: 1) swiveling is an intrinsic motion of RNAP; 2) the swiveling motion of RNAP (global conformational change) is directly linked to trigger loop folding (local conformational change); and 3) NTP binding at the active site influences the equilibrium of the swiveled (without NTP) and non-swiveled (with NTP) states of RNAP.

In the *Mtb* RNAP-*Mtb* NusG TEC without the RNA hairpin, we observed extra density extending from the C-terminus of the β subunit, allowing us to build a model with 19 additional amino acid residues at the C-terminus of the β subunit (from E1151 to N1169). This β subunit tail is positioned below the β-flap domain and fills a gap between the Zn-binding (ZBD) and dock domains of the β’ subunit, which makes the RNA exit channel narrower (**Fig. 7A and B left**). The β subunit tail is disordered in the PTC, which allows accommodation of the RNA hairpin in the RNA exit channel (**Fig. 7A middle and B right**). Sequence alignment of the C-terminus of the β subunit in bacterial RNAPs revealed this β subunit extension of ∼30-40 residues in *Mtb* and closely related phyla but not in *E. coli* and its closely related phyla (**Fig. 7C**).

## Discussion

Transcription pausing is a key mechanism for regulating gene expression in all organisms (43). The structural basis for transcription pausing is largely inferred from studies using the transcription system from *E. coli*, a model Gram-bacterium (1, 13, 16, 17, 25, 35). In this study, we provide the structural basis for NusG-dependent pausing in *B. subtilis* and *Mtb*, model Gram+ bacteria, by determining a series of cryo-EM structures of *Mtb* RNAP with bound NusG from either *B. subtilis* or *Mtb*. These structures include TECs, PTCs, and complexes in the process of escaping from the PTC (**Fig. 8**). We revealed how NusG interacts with the TTNTTT motif in the ntDNA within the paused transcription bubble. This interaction leads to rearrangement of the transcription bubble and disruption of proper base paring of the upstream DNA duplex in the PTC (**Fig. 3B and D**). Structure-guided biochemical investigation identified NusG residues that play important roles in transcription pausing (**Fig. S5**). Furthermore, fitting T residues into the narrow cleft between NusG and the β-lobe domain provides an explanation for the observed preference of T at these positions over other nucleotides. The presence of T bases over C in the TTNTTT motif is advantageous since the C-5 methyl of T provides a hydrophobic group to favor van der Waals interactions with NusG and RNAP. In previous studies, we sequentially mutated these nucleotides and showed that the last three Ts of the <u>T</u>TNT<u>TT</u> motif (-8 to -6, underlined) are critical for NusG-dependent pausing, and that these residues were protected from oxidation by KMnO_4_ in footprinting studies (30, 42). Consistent with the published biochemical data, the PTC structure showed the flipping of the T_-6_ base, which fits tightly into a cavity of NusG to form van der Waals interaction (**Fig. 3C**). Reduced pause half-lives and pausing efficiencies in the presences of alanine substituted NusG variants strengthen our structure-based interpretations (**Fig. S5**).

### The mechanism of NusG-dependent pausing

Cellular RNAP is a flexible and dynamic macromolecule, which changes its conformation throughout the transcription cycle. The DNA binding main channel of the bacterial RNAP core enzyme is widely opened, whereas it is closed during transcription elongation (44). The main channel is also closed when it binds a promoter recognition σ factor to form a holoenzyme (45). This type of RNAP motion is known as “clamp opening and closing”, which is a conserved motion in most cellular RNAPs (46). Swiveling is another motion of RNAP in which the swivel module rotates a few degrees (1∼4°) relative to the core module using a rotation axis roughly parallel to the bridge helix. The swiveling motion has been observed in PTCs formed with *E. coli* RNAP (17, 22, 41), and the elongation factors NusA and NusG influence the equilibrium between the swiveled and non-swiveled states of RNAP (41). In this study, we revealed that swiveling is an intrinsic motion of *Mtb* RNAP not only during the formation of PTC but also during transcription elongation, which is directly linked to the trigger loop motion (folding and unfolding) (**Figs. 5, 6 and 8, Movie S6**). NTP binding at the active site shifts the equilibrium from the swiveled to non-swiveled states, as well as the trigger loop from unfolded to folded states (with NTP). Since domains and motifs of RNAPs are highly conserved in all cellular RNAPs including bacterial, archaeal and three nuclear eukaryotic RNAPs (RNAPI, RNAPII and RNAPIII) (46–48), the coupling between swiveling and trigger loop folding may occur in all organisms.

RNAP swiveling changes the width of the cleft between NusG and the β-lobe domain of RNAP (**Figs. 4B and 8, Movie S6**). The same cleft is used for inserting the T residues of ntDNA during formation of the PTC, which restricts swiveling. Thus, during PTC formation, the ntDNA containing the TTNTTT motif acts as a wedge to prevent the transition from the swiveled to non-swiveled states. Since swiveling and trigger loop folding are linked, the NusG-ntDNA interaction can pause RNA synthesis despite it being ∼60 Å away from the active site of RNAP. Once the PTC is formed, removal of the T residues from the NusG/β-lobe cleft is required for pause escape such that RNAP can resume transcription elongation.

RNAP swiveling also changes the width of the RNA exit channel since it is positioned at an interface between the swivel and core modules of RNAP and is slightly narrower in the non-swiveled state compare with the swiveled state (**Fig. 7A**). Thus, RNA hairpin formation is only allowed in the PTC (**Fig. 5B**). The RNA hairpin is not essential for NusG-dependent pausing, but it increases the pause half-life. A wide range of pause strengths *in vivo* and pause half-lives *in vitro* were observed for various NusG-dependent pause sites (31). Our study provides mechanistic insight into how pause strength/half-life is determined; the strength of the NusG-ntDNA interaction (number of T residues in the transcription bubble and/or upstream DNA), together with the stability of the RNA hairpin, can determine the transition energy barrier required for RNAP to escape from the swiveled state. Higher energy barriers increase the pause strength/half-life.

### NusG is a transcription pausing factor in bacteria and the ntDNA in the transcription bubble provides a signal to influence transcription

NusG is a universally conserved transcription elongation factor that is present in all organisms (Spt5 in archaea and eukaryotes) and has been recognized as an anti-pausing factor and a core regulator of transcriptional polarity based on studies using the *E. coli* transcription system (49, 50). In *B. subtilis*, NusG stimulates pausing by establishing sequence-specific contacts between its NGN domain and conserved T residues of the ntDNA within the transcription bubble (30, 31). The structural and biochemical studies of NusG-dependent pausing presented here propose an alternative and perhaps more universally conserved view in which NusG functions as a pausing factor in bacteria. Multiple-sequence alignment analysis showed that most of the NusG residues involved in ntDNA interaction are conserved throughout the bacterial domain with exceptionally high conservation among firmicutes, actinobacteria and cyanobacteria (**Fig. S6**). Furthermore, we confirmed that *Mtb* NusG stimulates pausing of *Mtb* RNAP *in vitro* and found that pausing occurs at the same position as with *B. subtilis* RNAP and NusG (**Fig. 1D**). The conservation of pause promoting NusG sequences suggests that NusG might function as a transcription pausing factor in most bacteria. A key question that needs to answered is whether related bacteria also have critical TTNTTT motifs at specific positions that contact NusG to regulate transcription elongation.

The surface exposed single-stranded ntDNA in the transcription bubble within the TEC is one of a limited number of binding platforms for a transcription regulator to directly access nucleic acid when RNAP is bound and transcribing RNA. In addition to *B. subtilis* NusG, Spt5 in yeast (34) and the C128 subunit of RNAPIII (51, 52) regulate transcription elongation by contacting a T-rich ntDNA strand in the transcription bubble, suggesting that critically positioned polyT signals in ntDNA play an important role in transcription elongation and that the associated regulatory processes might be a universally conserved mechanism shared by all organisms including bacteria, archaea and eukaryotes.

## Materials and methods

### *Purification of Mtb* RNAP

The recombinant *Mtb* RNAP core enzyme (α_2_ββ’ω) was over-expressed in *E. coli* strain BL21(DE3)-CodonPlus-RIPL transformed by plasmid pMTBRP-5 (53). Overexpression of protein was induced by adding 1 mM isopropyl β-D-thiogalactopyranoside (IPTG) and grown at 19 °C for 22 h. Cell pellet was resuspended in 100 ml of lysis buffer (40 mM Tris-HCl, pH 7.9, 500 mM NaCl, 5% glycerol, 2 tablets of cOmplete EDTA-free protease inhibitor cocktail (Roche), 0.1 mg/ml lysozyme, and 1 mM β-mercaptoethanol). Cells were disrupted by sonication and the lysate was cleared by centrifugation (30,000 g for 20 min at 4 °C). The lysate was then loaded onto a 5 ml HisTrap HP column (Cytiva) equilibrated with buffer containing 40 mM Tris-HCl (pH 7.9), 500 mM NaCl, 5% glycerol, 1 mM β-mercaptoethanol, and 10 mM imidazole. After washing with buffer containing 30 mM imidazole, RNAP was eluted in buffer containing 40 mM Tris-HCl (pH 7.9), 100 mM NaCl, 5% glycerol, 0.2 mM β-mercaptoethanol, and 250 mM imidazole. Proteins were loaded onto a 1 ml heparin HP column (Cytiva) equilibrated with buffer A (20 mM Tris-HCl, pH 7.9, 5% glycerol, 0.2 mM β-mercaptoethanol, and 0.2 mM EDTA) plus 100 mM NaCl. After washing with 30% of buffer B (buffer B is 20 mM Tris-HCl, pH 7.9, 1 M NaCl, 5% glycerol, 0.2 mM β-mercaptoethanol, and 0.2 mM EDTA), RNAP was eluted in 60% of Buffer B. Proteins were concentrated using an ultracel-100 membrane filter unit (Millipore) and then the concentration of NaCl was adjusted to 100 mM. The resulting sample was then loaded onto a 1 ml HiTrapQ HP column (Cytiva) equilibrated in 10% of Buffer B. RNAP was eluted using a linear gradient from 10% to 80% Buffer B (20 column volumes). The fractions containing RNAP were pooled and concentrated to ∼10 mg/ml together with exchanging buffer (20 mM Tris-HCl, pH7.9, 200 mM NaCl, 5% glycerol, 0.2 mM β-mercaptoethanol, and 0.2 mM EDTA) and stored at -80 °C in small aliquots.

### *Purification of B. subtilis* RNAP

The *B. subtilis* RNAP core enzyme was purified from σ^A^ mutant strain ORB5853 [*rpoC*-His10 σ^A^(L366A)] (54), in which the L366A substitution in σ^A^ weakens the interaction between core enzyme and σ^A^. *B. subtilis* cells were grown in 2xYT (yeast extract/tryptone) liquid media containing 5 µg/ml chloramphenicol and 75 µg/ml neomycin at 37 °C until the OD_600_ of the culture reached 0.8. The pellet (30 g) was dissolved in 100 ml lysis buffer (50 mM Tris-HCl, pH 8.0, 500 mM NaCl, 10% glycerol, 2 mM β-mercaptoethanol and 5 mM MgCl_2_) containing 2 tablets of cOmplete EDTA-free protease inhibitor cocktail and 0.2 mg/ml lysozyme. Cells were disrupted by sonication and the lysate was cleared by centrifugation (30,000 g for 20 min at 4 °C). The lysate was then loaded onto a 5 ml HisTrap HP column (Cytiva) equilibrated with buffer containing 50 mM Tris-HCl (pH 8.0), 500 mM NaCl, 10% glycerol, and 2 mM β-mercaptoethanol. After washing with buffer containing 20 mM imidazole, the RNAP core enzyme was eluted in buffer containing 50 mM Tris-HCl (pH 8.0), 100 mM NaCl, 10% glycerol, 2 mM β-mercaptoethanol, and 200 mM imidazole. RNAP-containing samples were pooled and then loaded onto a 1 ml heparin HP column (Cytiva) equilibrated with buffer A (20 mM Tris-HCl, pH 7.9, 5% glycerol, 0.2 mM β-mercaptoethanol, and 0.2 mM EDTA) plus 100 mM NaCl. After washing with 30% of buffer B (buffer B is 20 mM Tris-HCl, pH 7.9, 1 M NaCl, 5% glycerol, 0.2 mM β-mercaptoethanol, and 0.2 mM EDTA), the RNAP core enzyme was eluted in 60% of Buffer B. RNAP containing samples were concentrated using an ultracel-100 membrane filter unit (Millipore) and then the concentration of NaCl was adjusted to 100 mM. The resulting sample was then loaded onto a 5 ml anion-exchange (Source 15Q; Cytiva) column equilibrated with buffer C (10 mM HEPES-KOH pH 8.0, 0.2 mM EDTA, 4 mM dithiothreitol (DTT)) plus 100 mM NaCl. *B. subtilis* RNAP was eluted using a linear gradient from 10% Buffer D (10 mM HEPES-KOH, pH 8.0, 0.2 mM EDTA, 4 mM DTT, and 1M NaCl) to 80% Buffer D (15 column volumes). The RNAP fractions were pooled and concentrated to 10 mg/ml and exchanged into a buffer containing 10 mM HEPES, pH 8.0, 200 mM NaCl, 5% glycerol, 0.5 mM DTT, and 0.2 mM EDTA) and stored at -80 °C in small aliquots.

### *Purification of B. subtilis* σ^A^

For production of *B. subtilis* σ^A^, *E. coli* BL21(DE3) cells were transformed with plasmid pSUMO-*Bsub*σ^A^ and then grown in 500 ml LB media supplemented with 50 μg/ml of kanamycin at 37 °C until OD_600_ = ∼0.5, at which time IPTG (0.4 mM final conc.) was added to overexpress σ^A^ by growing at 30 °C for 4 h. Cell pellet was resuspended in 50 ml lysis buffer (40 mM Tris-HCl, pH 7.9, 300 mM NaCl, 2 mM β-mercaptoethanol, and 5 % glycerol) supplemented with 1 tablet of cOmplete EDTA-free protease inhibitor cocktail and 0.1 mg/ml lysozyme. Cells were lysed by sonication and cleared by centrifugation (30,000 g at 4 °C for 20 min). Proteins were applied to a 1 ml HisTrap HP column (GE Healthcare), pre-equilibrated with buffer I (40 mM Tris-HCl, pH 7.9, 300 mM NaCl, 2 mM β-mercaptoethanol, and 5 % glycerol). The column was washed with buffer I containing 30 mM imidazole, and then eluted with buffer I plus 250 mM imidazole. σ^A^-containing fractions were pooled and dialyzed at 4 °C for 3 h with buffer I to remove imidazole. Proteins were subjected to Ulp1 protease digestion by incubating at 30 °C for 2 h to cleave the SUMO/His tags. Proteins were applied to a HiLoad 16/600 Superdex 200 gel filtration column (Cytiva) equilibrated with storage buffer (20 mM Tris-HCl, pH 7.9, 200 mM NaCl, 0.2 mM EDTA, 0.2 mM DTT, and 5 % glycerol). Fractions containing σ^A^ were pooled, concentrated with an Amicon ultra-4 centrifugation filter with a 10 kDa molecular weight cutoff (Merck Millipore), snap frozen and stored in -80 °C either directly (used for cryo-EM) or mixed with 100 % glycerol (50% final conc. for biochemical studies).

### Purification of NusG

For preparing the *B. subtilis* NusG, *E. coli* BL21(DE3) cells were transformed with plasmid pSUMO-*Bsub*NusG and grown in 500 ml LB media supplemented with 50 μg/ml of kanamycin at 37 °C to OD_600_ = ∼0.5. Overexpression was induced by adding 0.5 mM IPTG and grown at 30 °C for 4 h. Cell pellet was resuspended in 50 ml lysis buffer (40 mM Tris-HCl, pH 7.9, 300 mM NaCl, 2 mM β-mercaptoethanol, and 5% glycerol) supplemented with 1 tablet of cOmplete EDTA-free protease inhibitor cocktail and 0.1 mg/ml lysozyme. Cells were disrupted via sonication as described above and cleared by centrifugation. The cleared lysate was applied to a 1 ml HisTrap HP column (Cytiva), pre-equilibrated with buffer I (40 mM Tris-HCl, pH 7.9, 300 mM NaCl, 2 mM β-mercaptoethanol, and 5 % glycerol). The column was washed with buffer I containing 30 mM imidazole, and then eluted with buffer I containing 250 mM imidazole. NusG containing fractions were pooled and dialyzed at 4 °C for 3 h with buffer I to remove imidazole. The sample was then subjected to Ulp1 hydrolysis by incubating at 30 °C for 2 h to cleave the SUMO/His tags. Proteins were then applied to a HiLoad 16/600 superdex 75 gel filtration column (Cytiva) equilibrated with storage buffer (20 mM Tris-HCl, pH 7.9, 200 mM NaCl, 0.2 mM EDTA, 0.2 mM DTT, and 5 % glycerol). Fractions containing NusG were pooled, concentrated with an Amicon Ultra-4 centrifugal filter with a 5 kDa molecular weight cutoff (UFC8003, Merck Millipore), snap frozen and stored at -80 °C either directly (used for cryo-EM) or mixed 1:1 with 100 % glycerol (for biochemical studies). All derivatives of *B. subtilis* NusG and the wild-type *Mtb* NusG were purified similarly as described above.

### Promoter-initiated in vitro transcription pausing assay

Analysis of RNAP transcription pausing was performed as described previously with modifications (55). DNA templates were PCR-amplified from plasmids containing a strong promoter, a 29 nt C-less cassette, the predicted *coaA* pause hairpin, either a WT or mutant *coaA* TTNTTT pause motif, and downstream sequence. Halted elongation complexes containing a 29-nt transcript were formed by combining equal volumes of 2x template (100 nM) with 2x halted elongation complex master mix containing 80 µM ATP and GTP, 2 µM UTP, 100 µg/ml bovine serum albumin, 200 nM *B. subtilis* or *M. tuberculosis* RNAP with 400 nM *B. subtilis* σ^A^, 2 µCi of [α-^32^P]UTP and 2× transcription buffer (1x = 40 mM Tris-HCl, pH 8.0, 5 mM MgCl_2_, 5% trehalose, 0.3 mM EDTA, and 4 mM DTT). Reaction mixtures were then incubated at 37 °C for 5 min. Note that RNAP and σ^A^ were added from a 10x stock solution containing 0.75 mg/ml RNAP and 0.18 mg/ml σ^A^ in enzyme dilution buffer (1x = 20 mM Tris-HCl, pH 8.0, 40 mM KCl, 1 mM DTT, and 50 % glycerol). A 4x solution containing 4 µM NusG in 1x transcription buffer was then added, and the resulting reaction mixture was incubated at 23 °C for 5 min. For pausing assays, a 4x extension master mix containing 80 mM KCl, 600 µM of each NTP, 400 µg/ml rifampicin, in 1x transcription buffer was added, and the reaction was allowed to proceed at 23 °C. Aliquots of the transcription elongation reaction mixture were removed at various times and mixed with an equal volume of 2x stop/gel loading buffer (40 mM Tris-base, 20 mM Na_2_EDTA, 0.2% sodium dodecyl sulfate, 0.05% bromophenol blue, and 0.05% xylene cyanol in formamide). Transcripts were separated on standard 6% sequencing polyacrylamide gels, exposed to a phosphorscreen, and subjected to phosphorimaging using a typhoon phosphorimaging system (Cytiva). ImageJ software was used for quantification of transcripts on gel images. Pause half-life and pausing efficiency values were calculated by plotting relative intensities of the pause and run-off bands against the incubation time and fitting the data with a single exponential equation as described previously (30, 56).

### Preparations of the DNA-RNA scaffolds, PTC and TEC

To assemble RNA-DNA scaffolds, equimolar amounts of tDNA, ntDNA and RNA were mixed in annealing buffer (20 mM Tris-HCl, pH 8.0, 50 mM KCl, 5 mM MgCl_2_, 4 mM DTT, and 0.3 mM EDTA) followed by heating to 95 °C and then cooling to 10 °C (1.5 °C/min). The RNAs were 5’ end-labeled using polynucleotide kinase and γP^32^-ATP by incubating at 37 °C for 1 h. The upstream tDNA was not complementary to the RNA to prevent an extended RNA-DNA hybrid. Reconstitution of PTCs was performed by the addition of 200 nM RNAP core enzyme in 1x transcription buffer to 50 nM nucleic acid scaffolds, by incubation at 37 °C for 10 min. PTCs were transferred to ice and stored up to 2 h before use.

### In vitro transcription pause escape assays

Reactions were performed at 23 °C. PTCs and TECs containing 5′ end-labeled RNA were diluted in transcription buffer to 50 nM. NusG was then added to a final concentration of 1.7 μM. Rifampicin and KCl were added to final concentrations of 100 μg/ml and 50 mM, respectively. Extension of RNA was monitored following the addition of NTPs to the reaction. PTCs and TECs reconstituted with RNA A30 were elongated to C33 with an extension mix containing 5 μM GTP, 5 μM ATP, and 10 μM 3′-dCTP. Samples were removed at various times, quenched with an equal volume of 2x formamide loading buffer, and analyzed by denaturing 8% (19:1 acrylamide:bisacrylamide ratio) polyacrylamide gel electrophoresis.

### Sample preparation for Cryo-EM

*Mtb* RNAP (20 μM) and RNA-DNA hybrid (30 μM, either for PTC or TEC, **Fig. 2A**) were mixed in 1x transcription buffer (20 mM Tris-HCl, pH 8.0, 50 mM KCl, 5 mM MgCl_2_, 4 mM DTT, and 0.3 mM EDTA) at 37 °C for 10 min. NusG (30 μM, either from *B. subtilis* or *Mtb*) was added and further incubated for 10 min at 37 °C. 8 mM CHAPSO was added to the sample just before vitrification. When used, NTP was added to 1 mM final concentration.

### Cryo-EM grid preparation

Quantifoil grids (R 2/1 Cu 300 mesh, Electron Microscopy Sciences) were glow-discharged for 40 s prior to the application of 3.5 µl of the sample (∼6 mg/ml protein with 8 mM CHAPSO), and plunge-freezing in liquid ethane using a Vitrobot mark IV (FEI, Hillsboro, OR) with 100 % chamber humidity at 10 °C.

### Cryo-EM data acquisition and processing

Data was collected using a Titan Krios microscope (Thermo Fisher) equipped with a Falcon IV direct electron detector (Gatan) at the Penn State Cryo-EM Facility. Sample grids were imaged at 300 kV, with an intended defocus range of -2.5 to -0.75 μm with a magnification of 75,000x in electron counting mode (0.87 Å per pixel), and at a dose rate of 7.11 e^-^/Å^2^/second. Movies were collected with a total dose of 45 electrons per Å^2^. Movies were recorded with EPU software (Thermo Fisher Scientific) and data were processed by cryoSPARC3.3 (39). After motion correction and CTF estimation, 200 micrographs were used for picking particles (template-based autopicker followed by 2D-classification), and the selected particles were used for training the Topaz model (57). Particles were extracted with a 400-pixel box and subjected to multiple rounds of 2D classification to remove junk particles and subjected to multiple rounds of 3D heterogeneous refinements. The resulting best class was used for removing duplicated particles. CTF parameters were refined on a per micrograph and per particle basis using global and local CTF refinements, respectively. Particles were then subjected to a local motion correction followed by non-uniform refinement. The major variability component showed the RNAP swivel module rotation relative to the core module, coupled with the conformational changes of trigger loop and RNA hairpin. Particles were sorted into two or three clusters along the 3D conformational trajectories followed by 3D refinements (see cryo-EM pipelines of figures S1, S2, S7, S8 and S9 for details). Nominal resolutions were determined by using the goldstandard Fourier Shell Correlation (FSC) 0.143. The cryo-EM density maps were improved by Local anisotropic sharpening by Phenix (58).

### Model building, refinement, and analysis

A previously determined cryo-EM structure of *Mtb* RNAP (PDB: 5ZX2, removed SigE, DNA and RNA) (59) was fit into the cryo-EM maps using UCSF Chimera-v1.1 (60). Models of *B. subtilis* and *Mtb* NusG were made by Alphafold (61), and models of DNA and RNA were built using the *E. coli* RNAP TEC (PDB: 7MKO) as a guide, followed by fitting into the cryo-EM maps using Coot (62). Models were further improved using rigid-body (separated by core and swivel modules; β-lobe, β-protrusion, β-flap and β’-i1 domains; NusG, RNA-DNA scaffold, see Fig. S6) and real-space refinement by Phenix (58, 63) using sharpened maps by cryoSPARC-v3.3 (39). For preparing figures and movies, cryo-EM maps and molecules were visualized by using UCSF ChimeraX-v1.4 (64) and PyMOL-v2.4 (The PyMOL Molecular Graphics System, Schrödinger, LLC).

### Swivel module rotation analysis

First, two structures of RNAP (*e.g*., swiveled and non-swiveled states such as PTC and eTEC+NTP, respectively) were aligned using the RNAP core modules by PyMOL align command. Second, select their swivel modules, analyze a rotation axis and an angle of the swivel module of target structure (*e.g.,* PTC) relative to a reference structure (*e.g.,* eTEC+NTP) using a script Rotation Axis in PyMOL (draw_rotation_axis.py, https://pymolwiki.org/index.php/RotationAxis) (41).

## Acknowledgments

We thank George Garcia at the University of Michigan for the pMTBRP-5 plasmid. We also thank Carol Bator, Sung Hyun Cho and Mike Carnegie at the Penn State Cryo-EM facility for supporting cryo-EM data collection. We thank David Gilmour and Jean-Paul Armache at Penn State University, as well as Alexander Yakhnin and Mikhail Kashlev at the National Cancer Institute, for helpful discussions. We thank Albert Weixlbaumer at the Institut de Génétique et de Biologie Moléculaire et Cellulaire for providing the methodology for measuring the RNAP swivel module rotation. This research was supported by National Institutes of Health (NIH) Grant GM098399 (to P.B.) and GM131860 (to K.S.M.).

## Author contributions

K.S.M. and P.B. conceptualized the project. R.K.V. performed the experiments. M.Z.Q. participated in initial optimization of cryo-EM sample and grid preparation, screening, and data processing. K.S.M. and R.K.V. processed cryo-EM data, refined and analyzed the structures and prepared figures and tables. R.K.V., K.S.M. and P.B. conducted formal data analysis. K.S.M. and R.K.V. wrote the original draft manuscript and P.B. edited it. All authors reviewed, edited and approved the manuscript. K.S.M. and P.B. secured funding.

---

**Figure S1:**
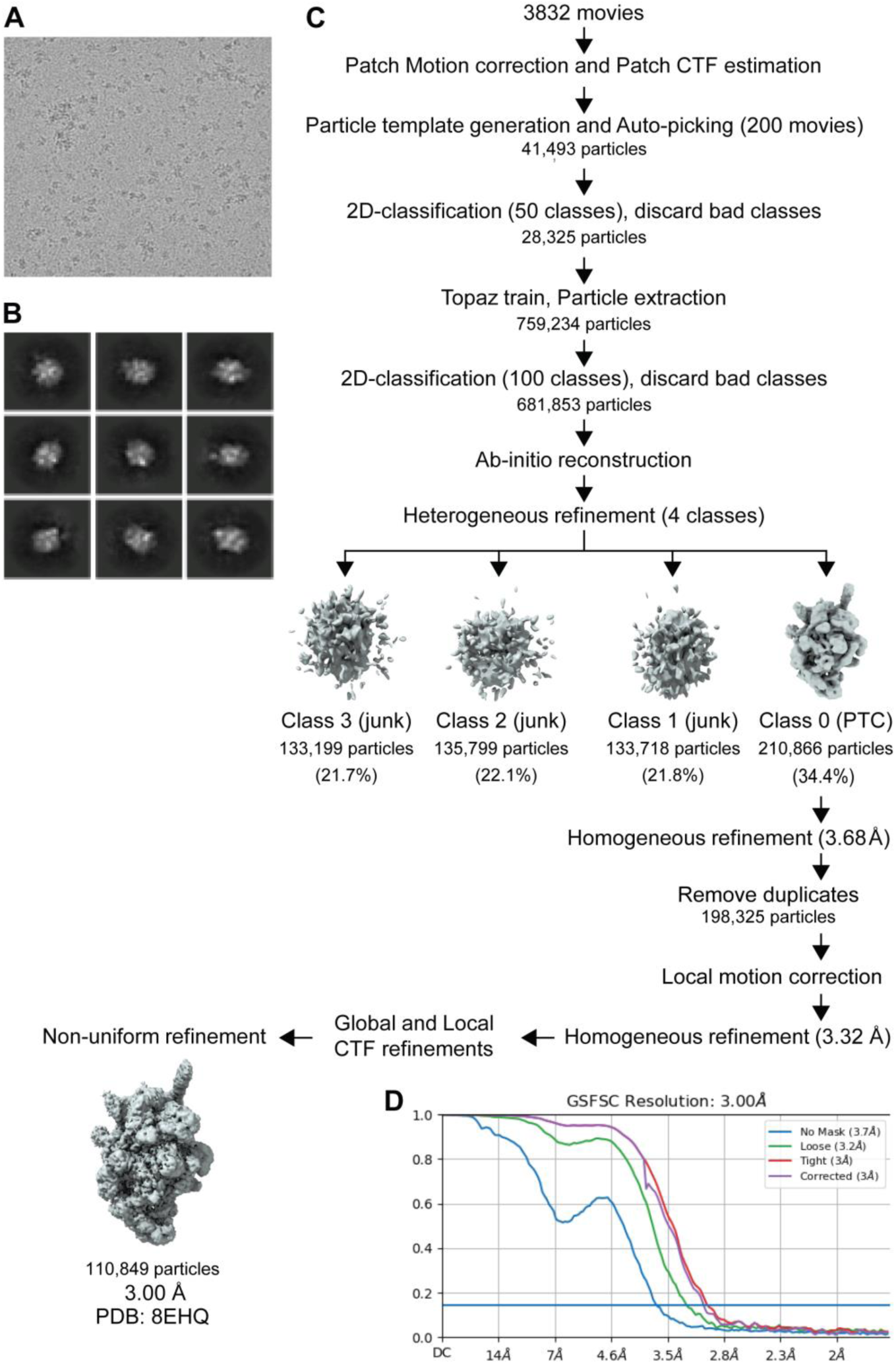
Cryo-EM data workflow for the PTC containing *Mtb* RNAP and *B. subtilis* NusG. **A.** Representative micrograph showing particle distribution. **B.** Selected representative 2D classes from 2D classification. **C.** Cryo-EM data processing and classification flow chart. **D.** Fourier shell correlation (FSC) plot for half-maps with 0.143 FSC criteria indicated. The nominal resolution was determined to be 3.00 Å.

**Figure S2:**
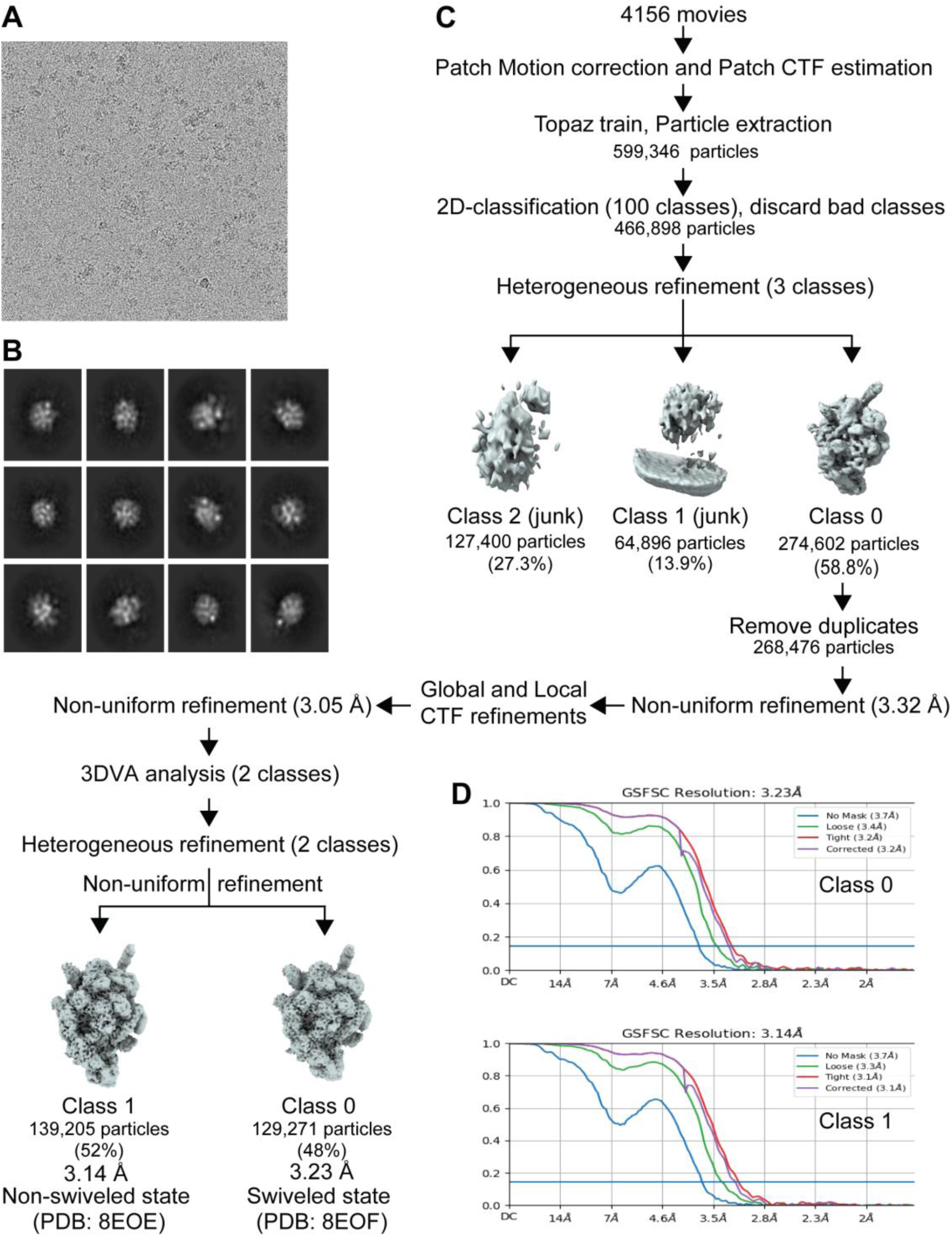
Cryo-EM workflow for the TEC containing *Mtb* RNAP and *B. subtilis* NusG. **A.** Representative micrograph showing particle distribution. **B.** Selected representative 2D classes from 2D classification. **C.** Cryo-EM data processing and classification flow chart. **D.** Fourier shell correlation (FSC) plot for half-maps with 0.143 FSC criteria indicated. 3DVA analysis resulted in two different structures with nominal resolutions of 3.23 Å (class 0) and 3.14 Å (class 1).

**Figure S3:**
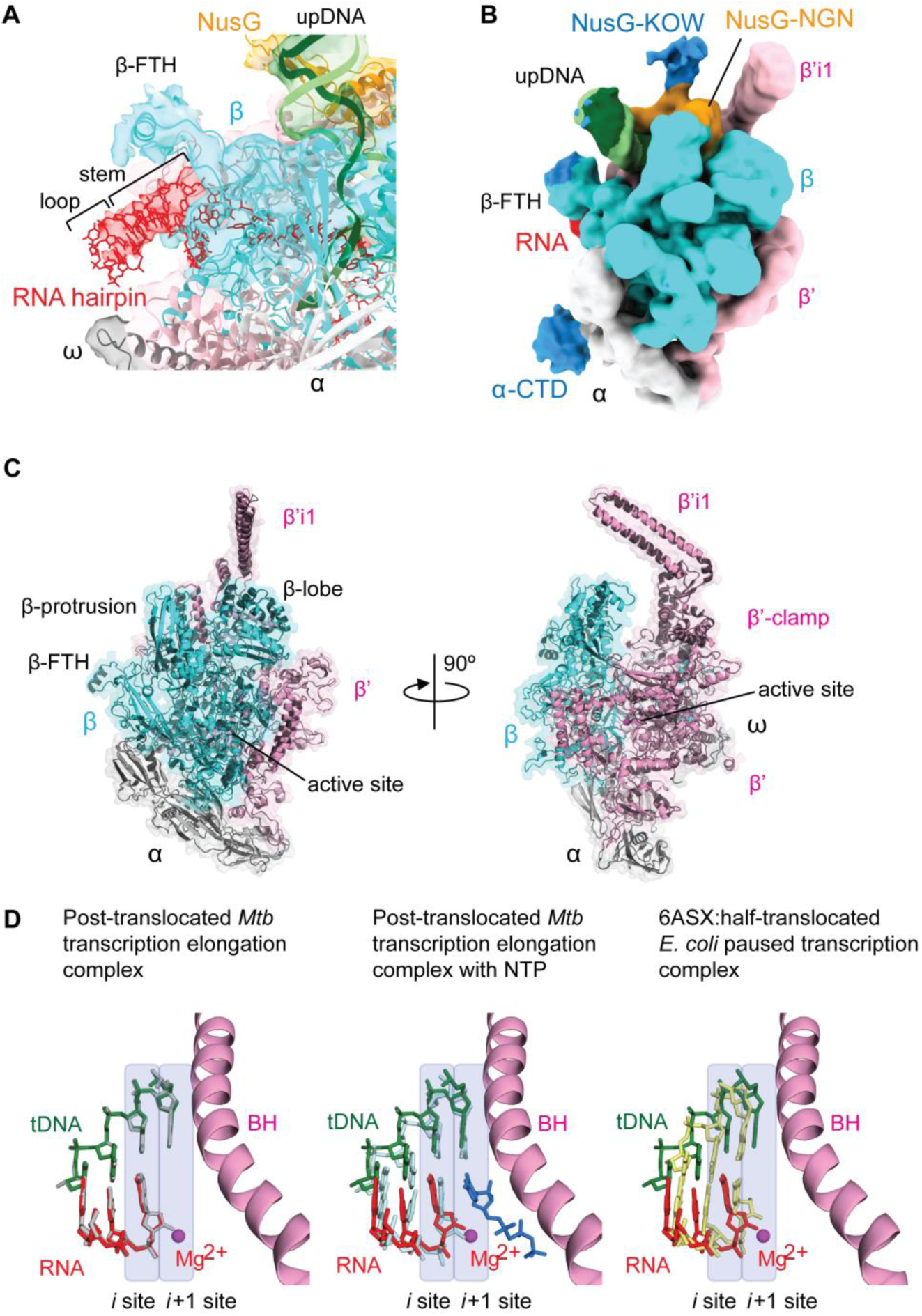
Cryo-EM structures of the PTC. **A.** Cryo-EM density map of the PTC centered at the RNA hairpin within the RNA exit channel. The transparent cryo-EM density map is colored according to the model in Fig. 3 (RNAP and DNA, ribbon models; RNA, stick model). Stem and loop of the RNA hairpin are indicated. β-FTH, flap tip helix. **B**. Low path filtered cryo-EM density map of the PTC shows the densities corresponding to the α-CTD and the NusG-KOW domains. β’i1, lineage-specific insertion. **C**. Comparison of the RNAP structures in the PTC (colored) and TEC (black). **D.** Comparison of the RNA-DNA hybrid (RNA, red; tDNA, green) proximal to the RNAP active site (active site Mg^2+^ ion and the bridge helix are shown as a magenta sphere and pink ribbon model, respectively) in the PTC with the TEC (left, gray), with the post-translocated TEC with NTP (middle, light blue and blue), and with the half-translocated *E. coli his* PTC (right, yellow). The active site (*i* and *i*+1 sites) is shown as transparent blue boxes.

**Figure S4:**
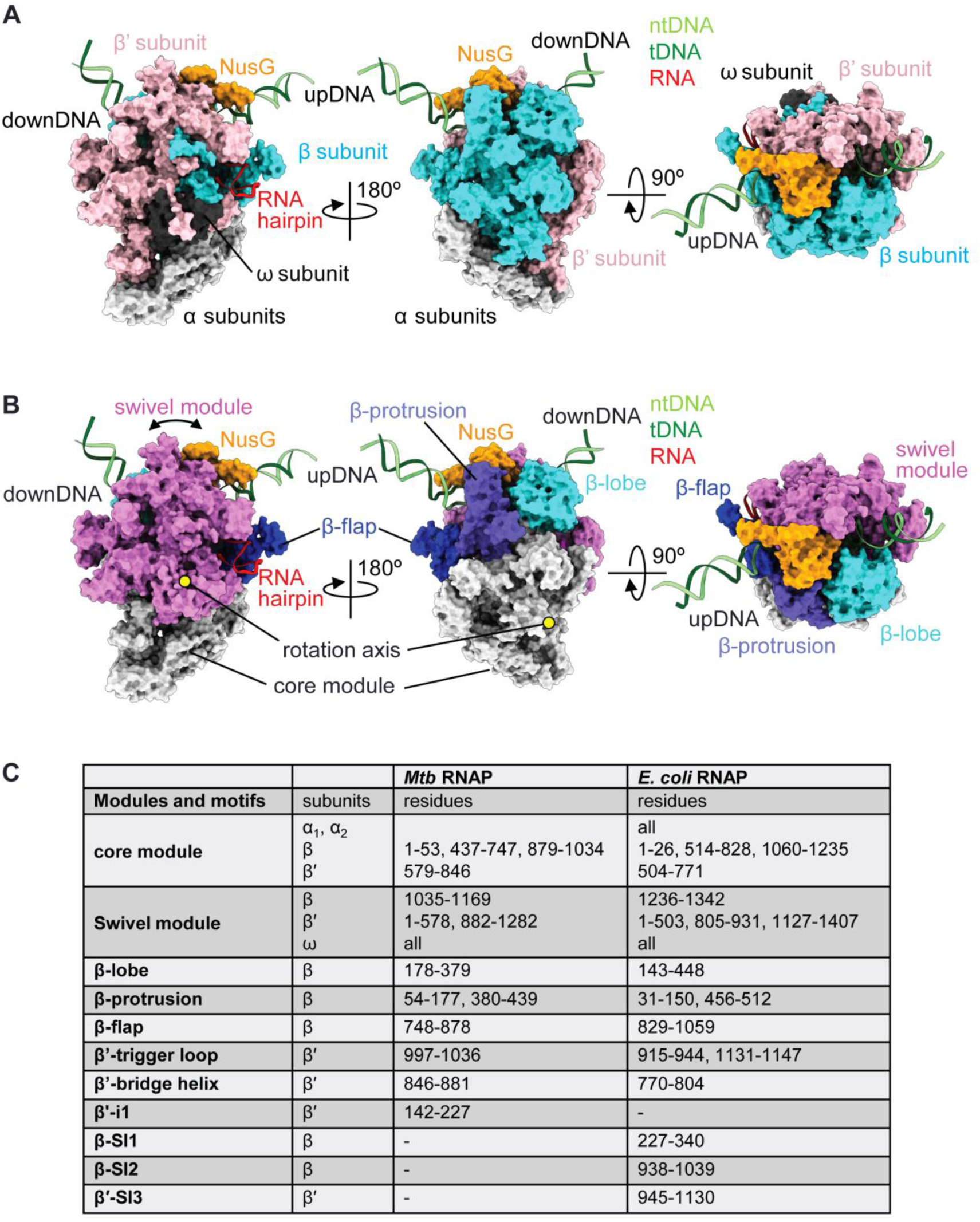
Structure of the *Mtb* RNAP PTC. Orthogonal views of the *Mtb* RNAP-*B. subtilis* NusG paused transcription complex indicating the subunits of RNAP **(A)** and the modules and domains of RNAP **(B)**. Location of the rotation axis of the swivel module is shown as yellow circles (left and middle). **C**. Modules and motifs of the *Mtb* and *E. coli* RNAPs.

**Figure S5:**
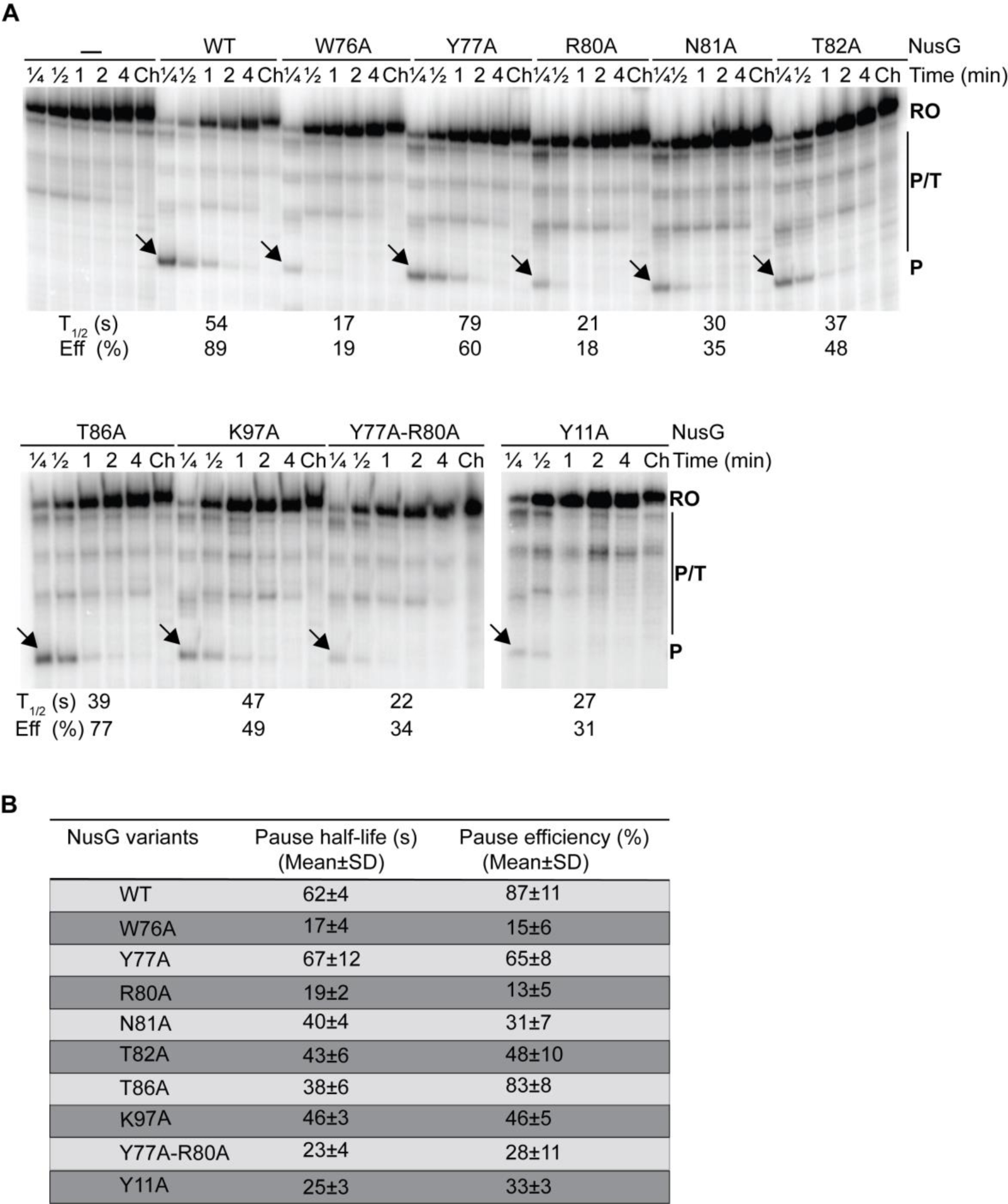
Structure-function analysis of NusG-dependent pausing. **A.** Gel image showing the effects of alanine substitutions on NusG-dependent pausing. Single-round *in vitro* transcription reactions were performed with the *coaA* template shown in Fig. 1A. Transcription was performed in the absence (–) and presence of 1 μM WT or mutant NusG as indicated. Reactions were stopped at the times shown above each lane. Chase reactions (Ch) were extended for an additional 10 min at 37 °C. Positions of paused (P) and run-off (RO) transcripts are marked. Additional pause, terminated, or arrested RNA species (P/T) were observed between P and RO. Pause half-life (t_½_) and pause efficiency (Eff (%)) values for this experiment are shown at the bottom of the gel. Although in the structure T82 of NusG did not show any interaction with the TTNTTT motif, the T82A variant was tested to corroborate the observation that a T82V variant caused pausing defects (1). **B.** Table showing RNAP pause half-life and pausing efficiency for WT and NusG mutants. Values are averages ± standard deviation (n=3).

**Figure S6:**
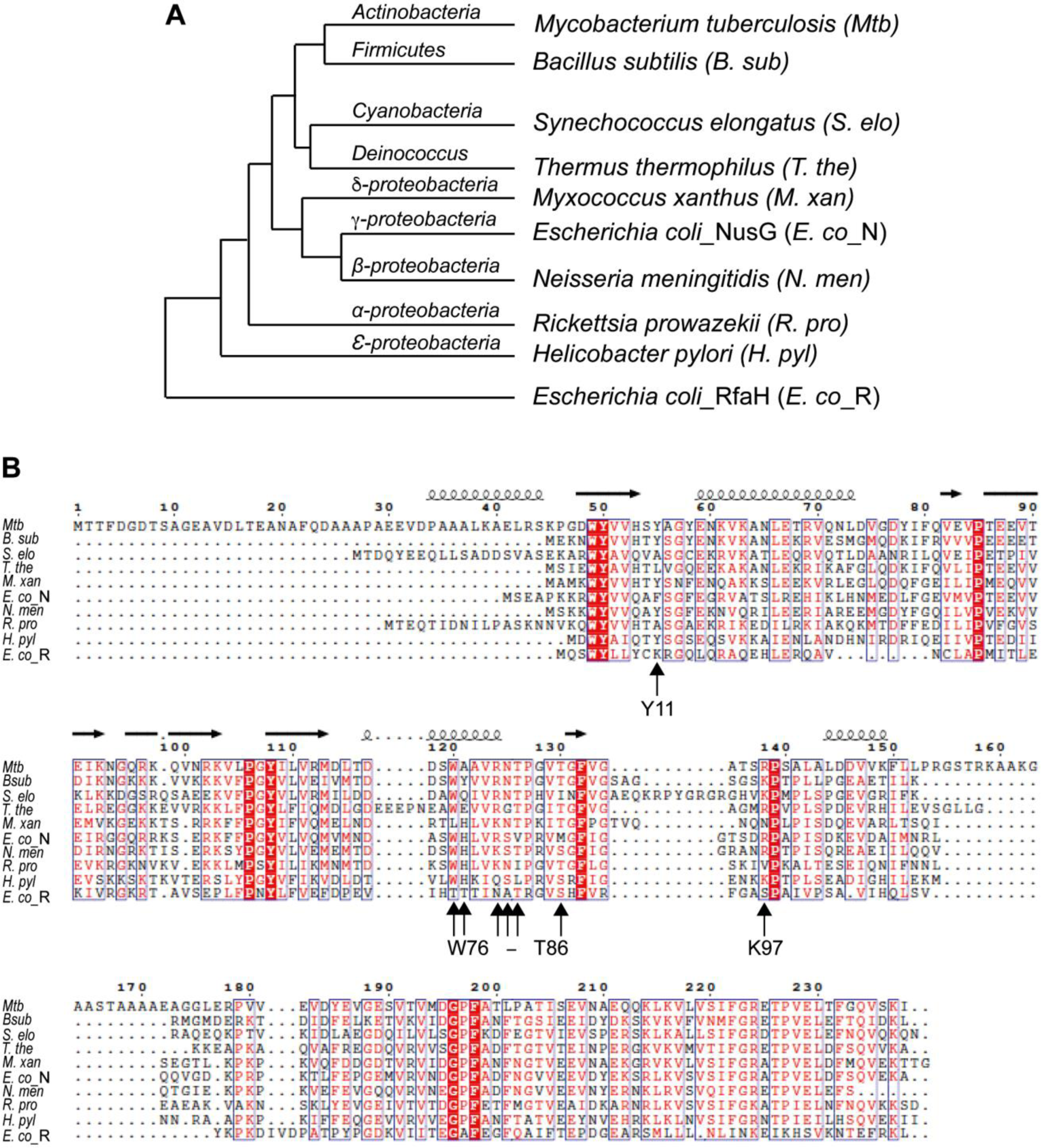
Phylogenetic analysis and amino acid sequence alignment of NusG. **A.** A phylogenetic tree was constructed using amino acid sequences of NusG from different bacteria including *E. coli* RfaH using the Neighbor-Joining method of the MEGA program (2)*. B. subtilis* and *M. tuberculosis* NusG clustered in the same branch. **B.** A sequence alignment (same sequences as used for phylogenetic analysis) using *Mtb* numbering was performed using the online Clustal-Omega program (3) and visualize online with ENDscript server (4). The boxed regions are conserved. Residues involved in interaction with ntDNA are marked with vertical arrows with the indicated *B. subtilis* numbering.

**Figure S7:**
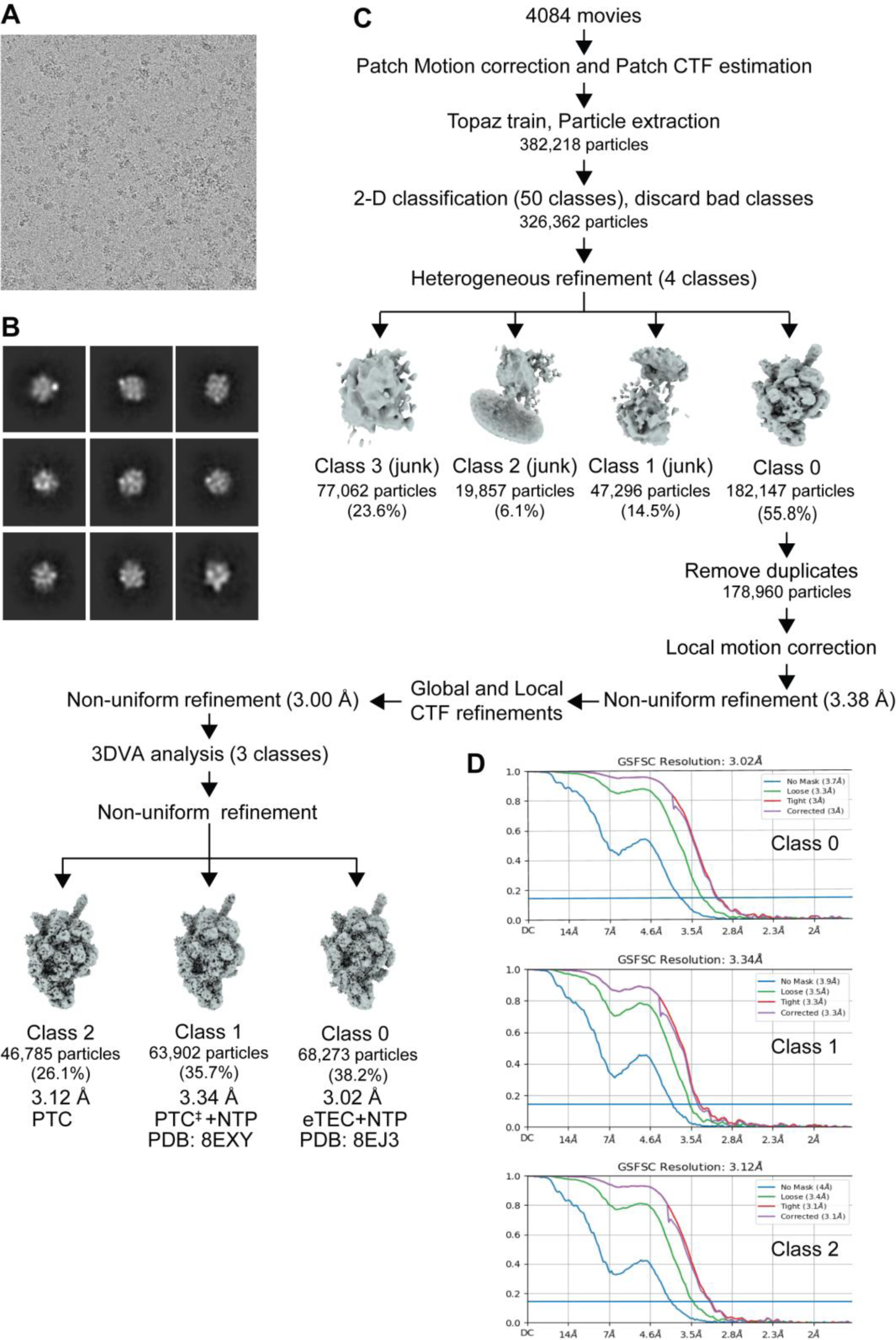
Cryo-EM workflow for the PTC, PTC^‡^ + NTP and eTEC + NTP. **A.** Representative micrograph showing particle distribution. **B.** Selected representative 2D classes from 2D classification. **C.** Cryo-EM data processing and classification flow chart. **D.** Fourier shell correlation (FSC) plot for half-maps with 0.143 FSC criteria indicated. 3DVA analysis resulted in three different structures: class 0 (3.02 Å), eTEC + NTP; class 1 (3.34 Å), PTC^‡^ + NTP; class 2 (3.12 Å), PTC. ^‡^ indicates that all of the structural features are identical to the PTC except for a slight rotation of the NusG/swivel module toward the downstream DNA and partial trigger loop folding.

**Figure S8:**
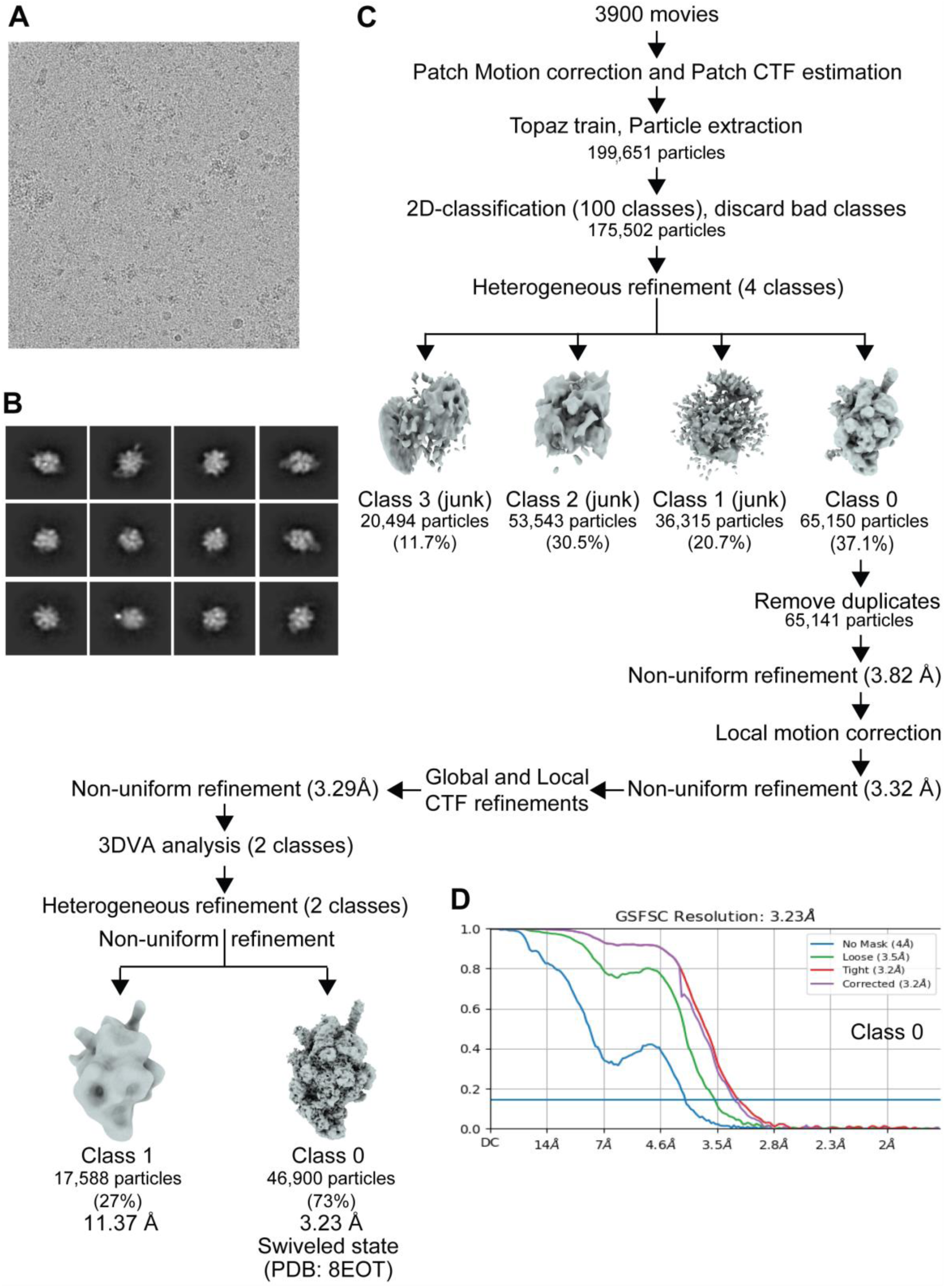
Cryo-EM workflow for the TEC containing *Mtb* RNAP and *Mtb* NusG. **A.** Representative micrograph showing particle distribution. **B.** Selected representative 2D classes from 2D classification. **C.** Cryo-EM data processing and classification flow chart. **D.** Fourier shell correlation (FSC) plot for half-maps with 0.143 FSC criteria indicated. 3DVA analysis resulted in two different structures with the majority of particles in class 0 having a nominal resolution of 3.23 Å (swiveled state).

**Figure S9:**
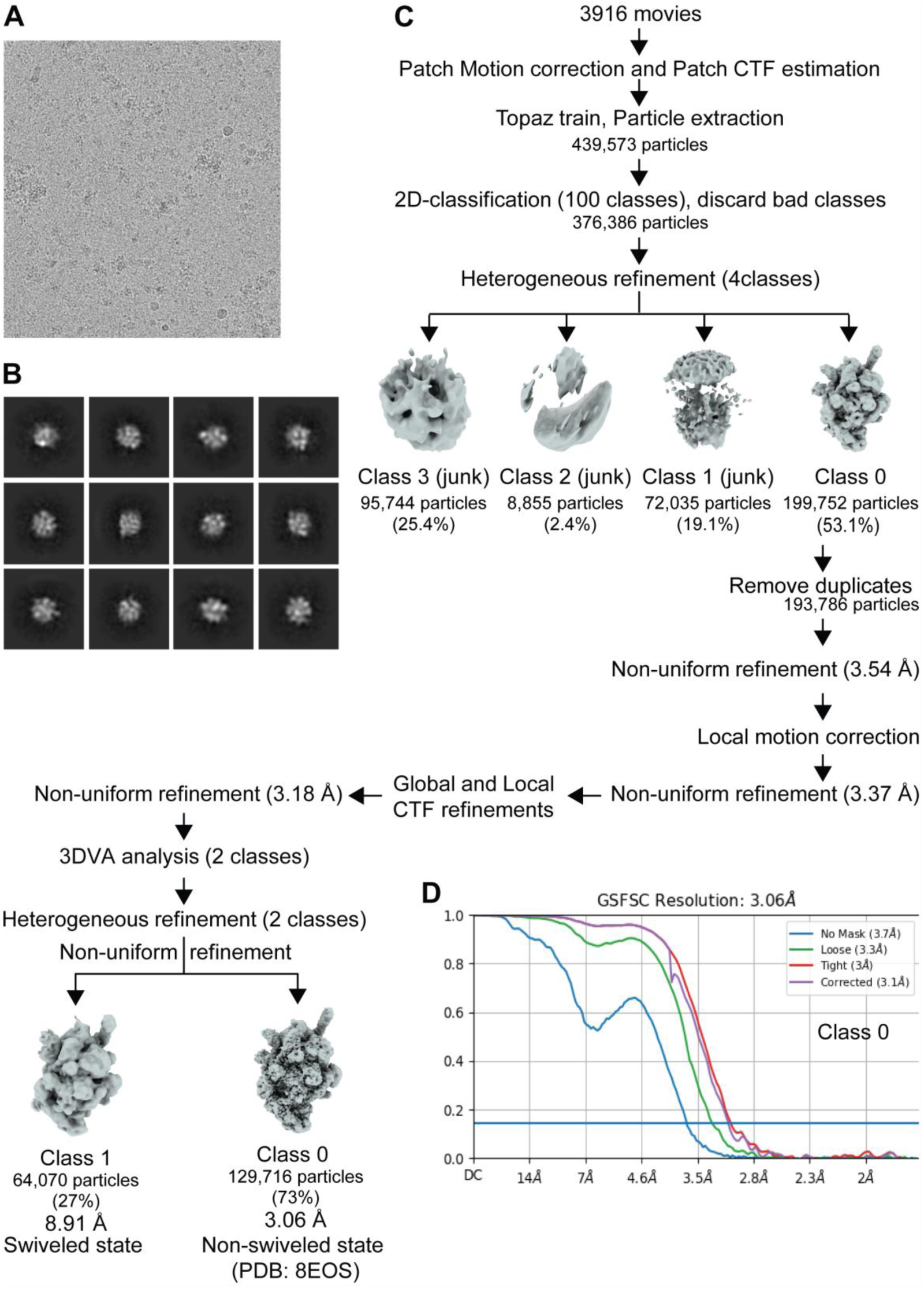
Cryo-EM workflow for the TEC containing *Mtb* RNAP and *Mtb* NusG with NTP. **A.** Representative micrograph showing particle distribution. **B.** Selected representative 2D classes from 2D classification. **C.** Cryo-EM data processing and classification flow chart. **D.** Fourier shell correlation (FSC) plot for half-maps with 0.143 FSC criteria indicated. 3DVA analysis resulted in two different structures with the majority of particles in class 0 having a nominal resolution of 3.06 Å (non-swiveled state).

## Legends for supplementary movies

**Movie S1: Cryo-EM density map of the NusG-dependent PTC.**

This movie shows the cryo-EM density map of the PTC in different angles. All of the components are colored and labeled as explained in the text.

**Movie S2: Structure of the NusG-dependent PTC.**

This movie highlights the PTC structural details. Swivel modules are indicated. For clarity, the lineage specific *Mtb* β’ insertion was removed from the structure.

**Movie S3: Principal component from 3D variability analysis (3DVA) of the cryo-EM data for the transcription complex with NusG.** This movie highlights the conformational changes of the swivel module of RNAP associated with changing the size of the RNA exit channel. Density maps of the first and last frames are light blue and light red, respectively, and intermediates are colored between light blue and light red. Modules, domains and the RNA exit channel are indicated before showing the RNAP motions.

**Movie S4: Principal component from 3D variability analysis (3DVA) of the cryo-EM data for the *Mtb* TEC with or without NTP.** This movie highlights the conformational change of the swivel module of RNAP. Density maps of the first and last frames are light blue and light red, respectively, and intermediates are colored between light blue and light red. Transcription complexes with (right) and without NTP (left) are shown side-by-side. Modules and domains are indicated before showing the RNAP motions.

**Movie S5: Conformational changes of *Mtb* and *E. coli* RNAPs.** Ribbon models of *Mtb* RNAP (left) and *E. coli* RNAP (right) of the TECs indicating the modules and domains. The movie highlights the conformational changes of RNAP from the non-swiveled to the swiveled states. Lineage specific insertions (β’-i1 in *Mtb* RNAP; β-Si1, β-Si2, β’-Si3 in *E. coli* RNAP) were removed for clarification. *Mtb* RNAP: non-swiveled state, TEC + NTP; swiveled-state, TEC without NTP. *E. coli* RNAP: non-swiveled state (PDB, 7Q0K); swiveled state (PDB, 7Q0J) (5).

**Movie S6: NusG-dependent pausing, and escape from the PTC.** The RNAP swivel module rotation (swiveling) directly links to nucleotide binding at the active site and to trigger loop folding. NusG-ntDNA interaction inhibits the transition from swiveled to non-swiveled states, thereby preventing trigger loop folding and RNA synthesis allosterically. Removal of the T residues from the NusG/β-lobe cleft is required for the rotation of swivel module and trigger loop folding for the RNAP continuing transcription elongation.

**Table S1.** Cryo-EM data collection, refinement, and validation statistics

|  | Paused transcription complex (PTC) | Transcription elongation complex (TEC) |  | PTC with NTP |  | TEC with *Mtb* NusG | TEC with *Mtb* NusG + NTP |
| --- | --- | --- | --- | --- | --- | --- | --- |
| **PDB ID** | 8EHQ | 8EOF | 8EOE | 8EXY | 8EJ3 | 8EOT | 8EOS |
| **EMDB ID** | 28149 | 28374 | 38373 | 28665 | 28174 | 28467 | 28466 |
| **RNAP conformation** | swiveled | swiveled | non-swiveled | swiveled (PTC + NTP) | non-swiveled (eTEC + NTP) | swiveled | non-swiveled |
| **Data Collection and processing** |  |  |  |  |  |  |  |
| Magnification | 75,000x |  |  |  |  |  |  |
| Voltage (kV) | 300 |  |  |  |  |  |  |
| Electron dose (e<sup>-</sup> / Å<sup>2</sup>) | 45 |  |  |  |  |  |  |
| Defocus range (μm) | -0.75 to -2.5 |  |  |  |  |  |  |
| Pixel size (Å) | 0.87 |  |  |  |  |  |  |
| Symmetry imposed | C1 |  |  |  |  |  |  |
| Initial particle images (no.) | 759,234 | 599,346 |  | 382,218 |  | 196,651 | 439,573 |
| Final particle images (no.) | 110,849 | 129,271 | 139,205 | 63,902 | 68,273 | 46,900 | 129,716 |
| Map resolution | 3.00 | 3.23 | 3.14 | 3.34 | 3.02 | 3.23 | 3.06 |
| (Å) |  |  |  |  |  |  |  |
| FSC threshold | 0.143 |  |  |  |  |  |  |
| Refinement |  |  |  |  |  |  |  |
| Initial model used (PDB code) | 6EDT | 8EHQ | 8EHQ | 8EHQ | 8EHQ, 8EOE | 8EOF | 8EOE |
| Model resolution (Å) | 3.00 | 3.30 | 3.20 | 3.34 | 3.02 | 3.30 | 3.10 |
| Model composition |  |  |  |  |  |  |  |
| Non-hydrogen atoms | 25,828 | 25,608 | 25,301 | 25,711 | 25,339 | 25,299 | 25,620 |
| Protein residues | 3,051 | 3,031 | 3,030 | 3,033 | 3,058 | 3,061 | 3,069 |
| DNA/RNA bases | 105 | 102 | 88 | 105 | 78 | 77 | 87 |
| Ligands | Zn:2, Mg:1 | Zn:2, Mg:1 | Zn:2, Mg:1 | Zn:2, Mg:1<br>G2P<br>(GMPCPP) | Zn:2, Mg:2<br>G2P<br>(GMPCPP) | Zn:2, Mg:1 | Zn:2, Mg:2<br>2TM (CMPCPP) |
| B factors (Å<sup>2</sup>) (mean) |  |  |  |  |  |  |  |
| Protein | 51.54 | 46.25 | 64.12 | 75.68 | 86.40 | 45.88 | 72.15 |
| DNA/RNA | 160.08 | 151.84 | 159.90 | 196.31 | 181.91 | 138.78 | 215.60 |
| Ligand | 87.95 | 78.82 | 88.13 | 71.02 | 83.80 | 78.23 | 62.31 |
| R.m.s. deviations |  |  |  |  |  |  |  |
| Bond lengths (Å) | 0.004 | 0.005 | 0.003 | 0.004 | 0.004 | 0.004 | 0.004 |
| Bond angles | 0.586 | 0.699 | 0.633 | 0.602 | 0.617 | 0.630 | 0.569 |
| (°) |  |  |  |  |  |  |  |
| **Validation** |  |  |  |  |  |  |  |
| **Mol Probity score** |  |  |  |  |  |  |  |
| Clash score | 5.42 | 5.21 | 5.34 | 5.39 | 5.38 | 5.09 | 20.12 |
| Rotamer outliers (%) | 6.46 | 17.45 | 17.38 | 20.98 | 15.79 | 6.03 | 3.59 |
| **Ramachandran plot** |  |  |  |  |  |  |  |
| Favored (%) | 95.36 | 89.68 | 90.41 | 89.79 | 89.91 | 94.42 | 94.66 |
| Allowed (%) | 4.58 | 9.55 | 9.06 | 9.45 | 9.20 | 5.45 | 5.18 |
| Disallowed (%) | 0.07 | 0.76 | 0.53 | 0.76 | 0.89 | 0.13 | 0.16 |
| **Model vs. Data** |  |  |  |  |  |  |  |
| CC (mask) | 0.87 | 0.87 | 0.84 | 0.86 | 0.86 | 0.87 | 0.85 |
| CC (box) | 0.74 | 0.78 | 0.72 | 0.79 | 0.70 | 0.78 | 0.68 |
| CC (peak) | 0.72 | 0.74 | 0.68 | 0.73 | 0.66 | 0.76 | 0.66 |
| CC (volume) | 0.84 | 0.83 | 0.80 | 0.83 | 0.83 | 0.84 | 0.82 |

